# Novel classes and evolutionary turnover of histone H2B variants in the mammalian germline

**DOI:** 10.1101/2021.09.05.459003

**Authors:** Pravrutha Raman, Callie Rominger, Janet M. Young, Antoine Molaro, Toshio Tsukiyama, Harmit S. Malik

**Author notes:** Correspondence to be addressed to: Harmit Singh Malik, 1100 Fairview Avenue N. A2-205, Seattle, WA 98109 USA.

## Abstract

Histones and their post-translational modifications facilitate diverse chromatin functions in eukaryotes. Core histones (H2A, H2B, H3, and H4) package genomes after DNA replication. In contrast, variant histones promote specialized chromatin functions, including DNA repair, genome stability, and epigenetic inheritance. Previous studies have identified only a few H2B variants in animals; their roles and evolutionary origins remain largely unknown. Here, using phylogenomic analyses, we reveal the presence of five H2B variants broadly present in mammalian genomes. In addition to three previously described variants (H2B.1, subH2B, and H2B.W), we identify and describe two new variants, H2B.L and H2B.N. Four of these five H2B variants originated in mammals, whereas H2B.L arose prior to the last common ancestor of bony vertebrates. We find that though mammalian H2B variants are subject to high gene turnover, most are broadly retained in mammals, including humans. Despite an overall signature of purifying selection, H2B variants evolve more rapidly than core H2B with considerable divergence in sequence and length. All five H2B variants are expressed in the germline. H2B.L and H2B.N are predominantly expressed in oocytes, an atypical expression site for mammalian histone variants. Our findings suggest that H2B variants likely encode potentially redundant but vital functions via unusual chromatin packaging or non-chromatin functions in mammalian germline cells. Our discovery of novel histone variants highlights the advantages of comprehensive phylogenomic analyses and provides unique opportunities to study how innovations in chromatin function evolve.

## Introduction

The genome and epigenome together determine form and function in all organisms. A significant component of the epigenome in eukaryotes is DNA-packaging units called nucleosomes. Eukaryotic nucleosomes typically contain ∼147bp of DNA spooled around an octamer of four core histones — H2A, H2B, H3, and H4 (Kornberg 1974; Kornberg and Thomas 1974; Thomas and Kornberg 1975; Luger, et al. 1997). These histone proteins share ancestry with histones from Archaea (Pereira, et al. 1997; Reeve, et al. 1997; Sandman and Reeve 2000; Ammar, et al. 2012; Mattiroli, et al. 2017; Talbert, et al. 2019) and some giant viruses (Erives 2017; Yoshikawa, et al. 2019; Liu, et al. 2021; Valencia-Sánchez, et al. 2021). All histone proteins possess a conserved histone fold domain (HFD) and more divergent N- and C-terminal tails. In eukaryotes, the core histones are typically expressed during genome replication with peak expression in S phase, to repackage genomes and their newly generated copies (Talbert and Henikoff 2021). Hence, they are also called replication-coupled (or RC) histones.

In addition to RC histones, eukaryotes encode histone variants to promote functional diversity and specificity in cellular processes. Histone variants are commonly expressed throughout the cell cycle. As a result, they are also referred to as replication-independent (or RI) histones. RI or variant histones replace RC histones in nucleosomes to promote specialized functions like DNA repair, chromosome segregation, and gene regulation (Talbert and Henikoff 2010; Martire and Banaszynski 2020). Typically, RC histones are present in eukaryotic genomes in large multi-copy arrays whereas histone variants are found either in one or a few copies. While histone variants resemble their RC histone counterparts, crucial differences in their HFDs distinguish their sequence and function. In addition, histone variants often differ more significantly from RC histones in their N- and C-terminal tails. These differences lead to deposition by different chaperones and distinct post-translational modifications, thus resulting in specialized functions by altering chromatin properties (Bönisch and Hake 2012; Henikoff and Smith 2015; Talbert and Henikoff 2017; Molaro, et al. 2018).

Some histone variants arose in early eukaryotic evolution, whereas others have evolved more recently in specific lineages (Malik and Henikoff 2003; Talbert and Henikoff 2010). Examples of ancient, well-conserved histone variants include H2A.Z, often found at transcription start sites, and CenH3, which localizes to centromeric DNA across most eukaryotes. However, other histone variants have evolved more recently in specific lineages (Eirín-López, et al. 2004; Yelagandula, et al. 2014; Rivera-Casas, et al. 2016; Molaro, et al. 2018). Many H2A variants, including macroH2A, H2A.W, and ‘short’ H2A variants, are found exclusively in filozoans (choanoflagellates and animals), plants, and placental mammals, respectively (Yelagandula, et al. 2014; Kawashima, et al. 2015; Rivera-Casas, et al. 2016; Molaro, et al. 2018). Although most eukaryotic histone variants appear to evolve under strong purifying selection, CenH3 also evolves adaptively in plant and animal lineages. CenH3’s rapid evolution has been proposed to be due to centromere drive or competition during female meiosis (Henikoff, et al. 2001; Malik and Henikoff 2009). Like ancient histone variants, most lineage-specific histone variants also evolve under strong purifying selection. However, some, like H2A.W in plants (Kawashima, et al. 2015) and short H2A variants in mammals (Molaro, et al. 2018), show signatures of adaptive evolution. Furthermore, while most histones are ubiquitously expressed, many lineage-specific histones, including some plant H2A.W variants and short H2A variants in mammals, are predominantly expressed in germ cells (Govin, et al. 2007; Boussouar, et al. 2008; Ferguson, et al. 2009; Molaro, et al. 2018; Khadka, et al. 2020; Lei and Berger 2020; Borg, et al. 2021). Such lineage-specific histone variants provide exciting opportunities to reveal novel epigenetic requirements and regulatory mechanisms via innovations in histone functions.

All four core RC histone proteins — H2A, H2B, H3, and H4 — are present in stoichiometric ratios within nucleosomes. However, there is non-uniform diversification of RC histones into histone variants. For example, there are many H2A variants in mammals but comparatively fewer H3 variants and even fewer H2B and H4 variants (Talbert and Henikoff 2010, 2021). This differential diversification may be due to each histone’s relative position in the nucleosome and its propensity to alter nucleosome properties upon replacement (Malik and Henikoff 2003). Yet H2B variants have proliferated in other lineages, including plants (Jiang, et al. 2020), suggesting they can be an abundant source of evolutionary and functional diversification. Our recent study revealed previously undescribed H2A variants and their evolutionary origins within mammals (Molaro, et al. 2018). Since H2A and H2B histones form obligate heterodimers, we investigated whether if there were H2B variants that remain undiscovered in mammalian genomes.

Three H2B variants — H2B.1, H2B.W, and subH2B— have been previous described in mammals, where they appear to be specialized for roles in the germline. H2B.1 (also referred to as testis-specific H2B, or TSH2B or TH2B) was one of the earliest H2B variants to be discovered from mammalian testes (Branson, et al. 1975; Shires, et al. 1975; Zalensky, et al. 2002). This variant is 85% identical in sequence to RC H2B and appears to play a role during spermatogenesis and in post-fertilization zygotes (Govin, et al. 2007; Montellier, et al. 2013; Shinagawa, et al. 2014). Structural and *in vitro* studies suggest that H2B.1-containing nucleosomes are less stable than RC H2B (Li, et al. 2005; Urahama, et al. 2014). This property has been suggested to facilitate histone-protamine exchange during spermatogenesis. More recently, H2B.1 has also been detected in mouse oocytes, where its function is not yet understood (Montellier, et al. 2013; Shinagawa, et al. 2014). H2B.W (also referred to as H2BFWT), was detected in sperm and appears to localize to telomeres when expressed in cultured cells (Churikov, et al. 2004; Boulard, et al. 2006); it remains functionally uncharacterized. SubH2B (for subacrosomal H2B), does not appear to function in chromatin and instead localizes to a perinuclear structure in sperm known as the subacrosome (Aul and Oko 2001). Although this compartment is involved in fertilization, subH2B’s exact function remains uncharacterized despite its abundant expression in sperm. In addition to these germline H2B variants, a fourth H2B variant, H2B.E, is expressed in olfactory neurons in rodent species. H2B.E differs from RC H2B by only five amino acid residues and plays important roles in regulating neuronal transcription and lifespan (Santoro and Dulac 2012). Preliminary evolutionary analyses of a few previously identified mammalian H2B variants (González-Romero, et al. 2010) suggested male germline-enriched variants may have accelerated rates of evolution. However, the evolutionary trajectories of H2B variants in mammals, including their diversity, origins and turnover, and their specialized germline functions are poorly understood.

Here, we perform detailed phylogenomic analyses of mammalian histone H2B variants and describe five evolutionarily distinct H2B variants in mammals, including two novel H2B variants, which we named H2B.L and H2B.N following previously proposed nomenclature guidelines (Talbert, et al. 2012). Except for H2B.L, which arose early in vertebrate evolution, all other H2B variants originated in early mammalian evolution and have been largely retained across mammalian orders. Yet, all H2B variants show dramatic expansions and/or pseudogenization, indicative of high evolutionary turnover. Whereas most H2B variants are predominantly expressed in testes or sperm, we find that the newly discovered H2B.L and H2B.N variants are instead overwhelmingly expressed in oocytes and early zygotes. Our analyses also reveal that H2B variants span a vast spectrum of evolutionary rates and have a wide range of sequence divergence from RC-H2B, suggesting that some variants might have evolved unconventional chromatin packaging properties or even non-histone functions. Together, our analyses reveal the presence of a larger H2B repertoire in mammals than was previously known, highlighting the power of evolutionary approaches to uncover innovation of lineage-specific chromatin functions.

## Results

### Seven distinct H2B variants in mammals

To identify variants of histone H2B in mammals, we interrogated genome assemblies from eighteen representative mammals. We performed comprehensive and iterative homology-based searches using both previously identified histone variants and new histone variants identified during our analyses (Molaro and Drinnenberg 2018) (see Methods). We further determined shared synteny (conserved genomic neighborhood) to identify orthologs. Thus, we were able to obtain a near-comprehensive list of all variant H2B open reading frames (ORFs) in these mammalian genomes. Since RC histones are present in large, nearly identical, multi-gene arrays, we did not compile all histone sequences that are near-identical to mouse or human RC H2B (Marzluff, et al. 2002; Talbert, et al. 2012). Although it is possible that some of those gene copies might be RI H2B variants, we focused instead on divergent H2B variants that are clearly distinct from RC H2B. Next, we performed protein sequence alignments of all identified H2B variants to identify incomplete sequences and manually curate our gene annotations. The alignment shows that H2B variants vary considerably in sequence and length in their N-terminal tails, making them difficult to align reliably in this region. Furthermore, we found that the C-terminal αC domain is absent or truncated in a subset of histone variants. Nevertheless, most H2B variants showed higher sequence conservation in their HFD and αC helix than in their tails (Figure 1B). To understand the evolutionary relationships between the H2B variant sequences we identified, we performed maximum likelihood phylogenetic analyses using PhyML (Guindon and Gascuel 2003; Guindon, et al. 2010). We used only regions we could reliably align across all variants, either an alignment of the HFD and αC domains (Figure 1A, Supplementary Figure S1) or just the HFD (Supplementary Figure S2). We did not observe any substantial differences between phylogenetic groupings or topology in our two analyses.

**Figure 1.**
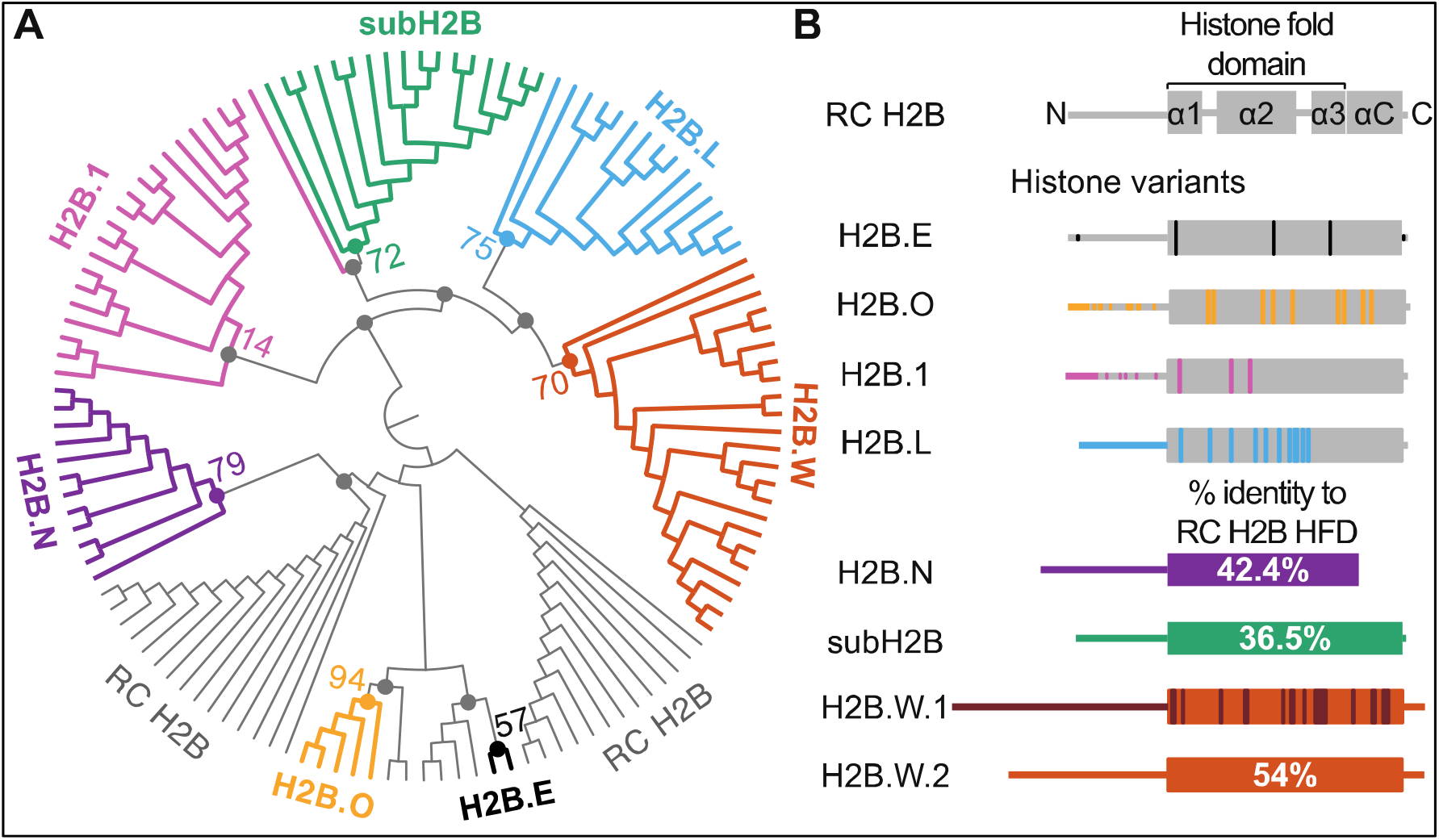
Phylogenomic analyses identify distinct H2B variant clades in mammals. (**A**) A maximum-likelihood protein phylogeny of the histone fold domain (HFD) of selected ancestral/replication-coupled (RC) H2B sequences and all intact H2B variants sequences from eighteen representative mammalian species is represented as a circular cladogram (see Supplementary Data 1 for a phylogram with branch lengths scaled to divergence). RC H2B histones are shown in grey, and seven H2B variant clades identified using phylogeny are highlighted in colors: H2B.E (black), H2B.O (yellow), H2B.N (purple), H2B.1 (pink), subH2B (green), H2B.L (blue), and H2B.W (orange). Bootstrap values at selected nodes with >50% support are shown along with colored dots to indicate the nodes they represent. The H2B.1 clade has a low bootstrap support of 14% owing to its high similarity to RC H2B (See Supplementary Figure S1, S3 for additional information). Select nodes with low bootstrap support values (<20%) are indicated with a grey dot. (**B**) Schematics of RC H2B and H2B variants. A structural schematic of a RC H2B at the top shows the N-terminus, histone fold domain or HFD (including the α1, α2, α3 helices and intervening loops), αC domain and the C-terminus. Variants with high identity to RC H2B (H2B.E, H2B.O, H2B.1 and H2B.L) are shown in grey with differences from RC H2B colored using the same colors as A. More divergent variants (H2B.N, subH2B and H2B.W.1 and H2B.W.2) are represented in solid colors and the percent identities of the HFD and αC domain compared to RC H2B are indicated. Differences between H2B.W.1 and H2B.W.2 are further highlighted in brown to indicate the divergence of these paralogs. Schematics and percent identities are based on human sequences, except for H2B.E and H2B.O, which are only found in rodents (mouse sequence used) and platypus respectively and subH2B which is pseudogenized in humans, (rhesus macaque sequence used).

Our analyses revealed seven distinct clades that represent discrete classes of H2B variants with unique features (Figure 1A, Supplementary Figure S1). Five of these clades are broadly distributed among mammals, whereas two clades have a restricted species distribution, suggestive of very recent evolutionary origins. The first of these species-restricted clades is *H2B*.*E*, which was originally identified through functional analyses of olfactory neurons in mice (Santoro and Dulac 2012) (Figure 1). We could only find one unambiguous ortholog of *H2B*.*E* in the closely-related rat genome but none in the more distantly-related guinea pig genome (Supplementary Figure S1). Since H2B.E differs from most copies of RC H2B by only five amino acid residues (three within the HFD) (Santoro and Dulac 2012), we recognized that phylogenetic analyses alone may not be adequate to identify all *H2B*.*E* orthologs. Therefore, we turned to shared synteny analyses to search for *H2B*.*E* orthologs in other mammalian genomes (Supplementary Figure S3A). Mouse *H2B*.*E* is found within a small cluster of *RC H2B* genes that is distinct from the major *H2B* cluster (Wang, et al. 1996; Marzluff, et al. 2002). Aligning all *H2B* genes within the *H2B*.*E* syntenic locus, we found only a single copy of *H2B* in mouse and rat that shares a majority of the five distinct residues characteristic of the originally identified mouse *H2B*.*E* (Supplementary Figure S3B). Based on this evidence, either *H2B*.*E* arose only in the rodent lineage, or *H2B*.*E* does not have enough distinguishing sequence features for us to unambiguously identify orthologs among the many closely related *H2B* genes present in mammalian genomes. Given this uncertain status, we do not further discuss *H2B*.*E* in our study.

Similarly, we identified a previously undescribed clade of H2B variants that we named *H2B*.*O* (we follow the histone nomenclature guidelines proposed in (Talbert, et al. 2012)), which is exclusively found in the platypus genome (Supplementary Figure S1). *H2B*.*O* variants represent a *bona fide* clade *i*.*e*., they group together to the exclusion of all other H2Bs. Their expression appears to be enriched in platypus’ germline tissues (testes or ovaries) albeit at low levels (Supplementary Figure S4). Due to their clear absence from the other mammals, we are unable to draw more significant conclusions about their function and evolutionary constraints.

Of the remaining broadly distributed H2B variants, three (H2B.1, subH2B, and H2B.W) have been previously described, whereas two (H2B.L, H2B.N) are newly identified by our analysis. Each of the three variants displayed unique evolutionary features. Most of the seven clades have high bootstrap support for the grouping of their orthologs (>50%) to the exclusion of other H2Bs. The only exception was that H2B.1 orthologs grouped together with low confidence (14%), likely due to their very high sequence similarity to RC H2B within the HFD used to generate the phylogeny (Figure 1B, Supplementary Figure S1). Although the N-terminal tail of most H2B variants is too diverged for use in phylogenetic analysis, H2B.1’s N-terminal tail can be reliably aligned to RC H2B. A phylogeny using a full-length alignment (*i*.*e*., including the N-terminal tail) unambiguously distinguished all H2B.1 orthologs from RC H2B with high bootstrap support (100%, Supplementary Figure S5). Synteny further supports unambiguous orthology for all the H2B.1 genes we examined (see below).

The H2B.W clade is also broadly present across mammals (Supplementary Figure S1). However, we also found that human *H2B*.*M*, which is found in genomic proximity to human *H2B*.*W*, groups within the H2B.W clade (Supplementary Figures S1, S7). Most features that distinguish human H2B.M from human H2B.W lie in the divergent N-terminal tails whereas their HFDs are much more similar (Figure 1B). We found many such apparent duplications of *H2B*.*W* histone variants across mammalian species. Their presence on the X chromosome and their close proximity to each other could allow copies to recombine or undergo gene conversion, resulting in similar sequences. We performed GARD analyses to test for such signatures of gene conversion and found that mammalian *H2B*.*W* variants are indeed undergoing recurrent gene conversion with each other, leading to the species-specific clustering pattern (Supplementary Figure S1). Based on the established guidelines for histone nomenclature (Talbert, et al. 2012), we henceforth refer to this clade as H2B.W in mammals; we refer to human *H2B*.*W* as *H2B*.*W*.*1* and *H2B*.*M* as *H2B*.*W*.*2*.

Like H2B.1 and H2B.W, we found that the subH2B clade is broadly represented across mammals (Supplementary Figure S1) except in human, where *subH2B* appears to be a pseudogene. We also found two phylogenetically distinct clades — H2B.L and H2B.N — that have not been previously identified. An unusual feature of both H2B.L and H2B.N is that they are encoded by intron-containing genes, whereas all other H2B variants and RC H2B lack introns (Supplementary Table S1). The intron in these two variants is in the same location with respect to the histone fold domain, suggesting that *H2B*.*L* and *H2B*.*N* may have a common ancestor, although our current phylogeny does not provide adequate support for their common origin.

Thus, our phylogenetic analyses identified seven distinct clades of H2B variants, including five that are broadly distributed among mammals. Although the H2B variant clades are clearly distinct from each other, and well supported by high bootstrap values, we are unable to make any strong inferences about the branching order of the clades *i*.*e*., whether they arose from a single duplication from ancestral *RC H2B* and subsequently diversified (monophyletic) or whether they arose via independent duplications of RC H2B (polyphyletic). This poor resolution contrasts with the strong evidence for monophyly of the short histone H2A variants in mammals (Molaro, et al. 2018).

### Structural features of H2B variants

To identify key residues that distinguish RC H2B from H2B variants, we compared the HFD and αC of RC H2B to each of the five broadly retained H2B variants. We chose orthologs from seven mammals, all of which encode at least one copy of all H2B variants (Figure 2A). We aligned orthologs of RC H2B and H2B variants and created logo plots to visualize their sequence conservation (Methods). To investigate each variant’s divergence from RC H2B, we calculated the Jensen-Shannon distance (JSD) at each position, comparing a set of 7 orthologs of each variant to a set of 7 orthologs of RC H2B from the same species. High JSD values indicate between-paralog differences that are also conserved within both groups of orthologs. We did not identify any residues that are conserved across all H2B variants, but different from RC H2B. However, we identified residues that are conserved across orthologs of each H2B variant but distinct from RC H2B (high JSD, >0.75, Figure 2A). We mapped these variant-specific conserved residues onto homology models constructed using a previously described structure of human RC H2B (PDB:5y0c, Figure 2B) (Arimura, et al. 2018).

**Figure 2.**
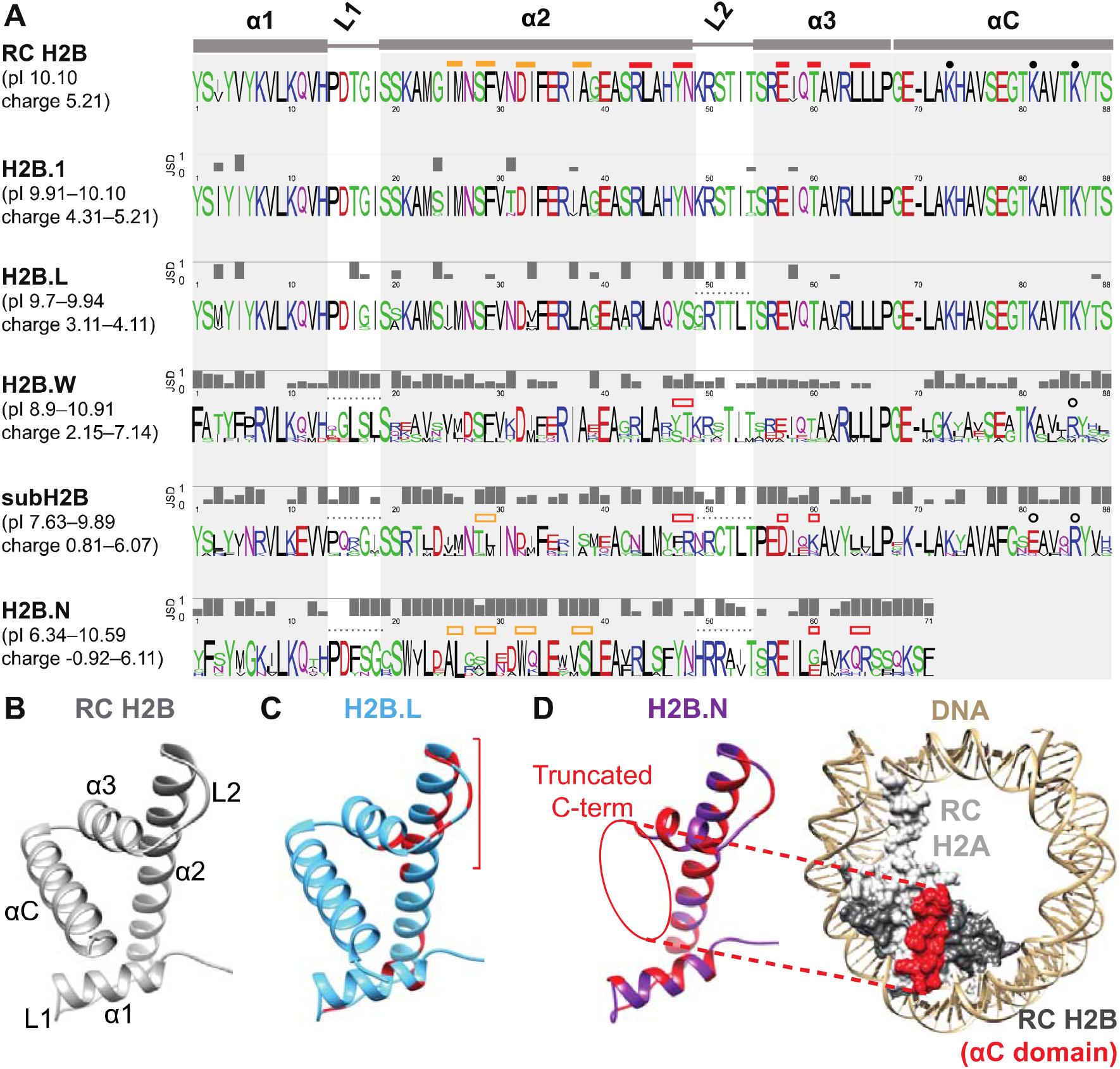
H2B variants diverge from RC H2B and each other in many protein features. (**A**) Logo plots depicting protein alignments of the HFD (α1, L1, α2, L2, α3) and αC domain of RC H2B and H2B variants across an identical set of representative mammals (see Methods). Colors of residues highlight their biochemical properties: hydrophobic (black), positively charged (blue), negatively charged (red), polar (green) and others (magenta). Jensen-Shannon distances (JSD) at each amino acid position were calculated between RC H2B and each H2B variant. Low JSD values indicate low divergence between RC H2B and variant H2Bs, whereas high JSD values indicates that RC H2B and H2B variants have distinct residues. Above the RC H2B logo, we indicate residues that interact with H2A (filled yellow boxes) or H4 (filled red boxes) and residues that are post-translationally modified (filled circles) (Luger, et al. 1997; McGinty and Tan 2021). Changes at these positions in H2B variants are indicated above each logo plot with empty boxes/circles, and loop regions with altered residues are indicated with a dotted line. The ranges of isoelectric points (pI) and charge across orthologs of each variant are shown in parentheses on the left. Note that H2B.N orthologs are missing most of the αC domain. (**B**) Structure of the HFD and αC domain of human RC H2B (Arimura, et al. 2018). The N and C terminus are unstructured. (**C**) Homology model of human H2B.L (blue) indicates a cluster of red sites (bracket) that differ from RC H2B (see Methods). (**D**) A homology model of human H2B.N (left, purple) with sites that differ from RC H2B highlighted in red (see Methods). RC H2A-H2B within a nucleosome (PDB:5y0c; (Arimura, et al. 2018)) is shown on the right, with DNA (tan), RC H2A (light grey), and RC H2B (dark grey) surfaces and the RC H2B αC domain (red) highlighted.

We find that H2B.1 and H2B.L are highly similar to RC H2B in their HFD (Figure 1B, 2A). H2B.1 differs from RC H2B by only 3 conserved differences in the HFD (JSD ∼1.0) (Figure 2A, Supplementary Figure 6A). The N-terminal tail of H2B.1 differs more from RC H2B (Supplementary Figure 6B), including at S/T residues that can be phosphorylated in RC H2B (Zalensky, et al. 2002; Li, et al. 2005). With this exception, all residues important for H2A (yellow bars), H4 (red bars) or DNA interaction and PTM residues (black dots) are identical to RC H2B, suggesting that H2B.1 and RC H2B share similar protein and chromatin properties.

Similarly, H2B.L orthologs have only a few fixed differences from RC H2B within the H2A-, H4-interacting residues and PTM sites, suggesting that these properties are likely conserved between RC H2B and H2B.L. Instead, most of the changes that distinguish H2B.L from RC H2B occur in other HFD sites, with several clustered around the second DNA binding loop (L2, Figure 2A, C) that could affect DNA binding or specificity. Furthermore, H2B.L is predicted to have a slightly lower charge than RC H2B (Figure 2A) that might result in less tightly-packed DNA. In contrast to its HFD, H2B.L’s N-terminal tail differs dramatically from RC H2B (Supplementary Figure 6B). For example, H2B.L’s N-terminal tail is missing key lysine residues that are post-translationally modified in RC H2B. Atypically for H2B proteins, H2B.L has a variable-length polyglutamine tract in its N-terminal tail that could facilitate protein-protein interactions (Schaefer, et al. 2012). Since H2B.L is a newly identified histone variant, its biochemical properties remain uncharacterized.

The remaining H2B variants (H2B.W, subH2B, H2B.N) share less than 50% amino acid identity with RC H2B in their HFD. For example, many residues in the HFD of H2B.W are conserved among orthologs, but diverged from RC H2B (JSD∼1, Figure 2A); these appear to cluster in the homology model (Supplementary Figure 6A). In spite of these differences, most H2A- and H4-interacting residues and PTM sites are conserved between H2B.W and RC H2B (Figure 2A). Divergence is even greater in their N-terminal tails, which cannot be reliably aligned with RC H2B (Supplementary Figure 6B). Human H2B.W.1 and H2B.W.2 also diverge most from each other in their N-terminal tail (Figure 1B). Unlike all other H2B variants, H2B.W variants have an extended C-terminal tail in some mammals (including humans), which is nearly identical between H2B.W.1 and H2B.W.2 in primates. Their divergence results in a wide range of charge and isoelectric points within H2B.W variants.

Even though subH2B localizes to the subacrosome in sperm (Aul and Oko 2001), it is nonetheless able to localize to chromatin in the nucleus when expressed in cell lines (Tran, et al. 2012). Putative H4-interacting residues, L2 residues and PTM residues are different between subH2B and RC H2B (open red/yellow boxes, Figure 2A, Supplementary Figure 6A, B). These differences may contribute to its unusual biological role outside the nucleus.

Finally, H2B.N shows the most dramatic differences from RC H2B in the HFD (Figure 2A). Although H2A-, H4-interacting residues and residues in L2 are largely conserved between H2B.N orthologs, they are highly divergent from RC H2B. The most striking difference is that most H2B.N orthologs are significantly truncated in their C-terminus.

Homology modeling predicts that this truncation results in the loss of the αC domain, whose residues are part of the essential nucleosome acidic patch (Nacev, et al. 2019; McGinty and Tan 2021) (Figure 2D). This suggests that the unusual H2B.N could endow nucleosomes with unique properties, or that H2B.N might have evolved non-nucleosomal functions like subH2B.

### Evolutionary origins of mammalian H2B variants

To identify the age and subsequent evolutionary patterns of the five mammal-wide clades of H2B variants, we searched genome assemblies of representative mammals and an outgroup, chicken. We classified uninterrupted open reading frames (ORFs) as intact genes (Figure 3A). We made the distinction between pseudogenes with many frame-disrupting mutations versus those that are only a single point mutation away from encoding an intact ORF (indicated with an asterisk); the latter could represent sequencing errors in otherwise intact ORFs. We found that all H2B variants have been largely retained across mammals at their shared syntenic location (Figure 3B, Supplementary Figure S7). Based on their presence in all eutherian (placental) mammals, we infer that both *H2B*.*1* and *H2B*.*W* clades arose in the last common ancestor of eutherian mammals (∼105MYA), whereas the *subH2B* and *H2B*.*N* clades also contain marsupial and platypus (but not chicken) sequences, and therefore arose in the last common ancestor of all mammals (∼177MYA) (Figure 3A) (divergence times calculated using TimeTree estimates (Hedges, et al. 2015)).

**Figure 3.**
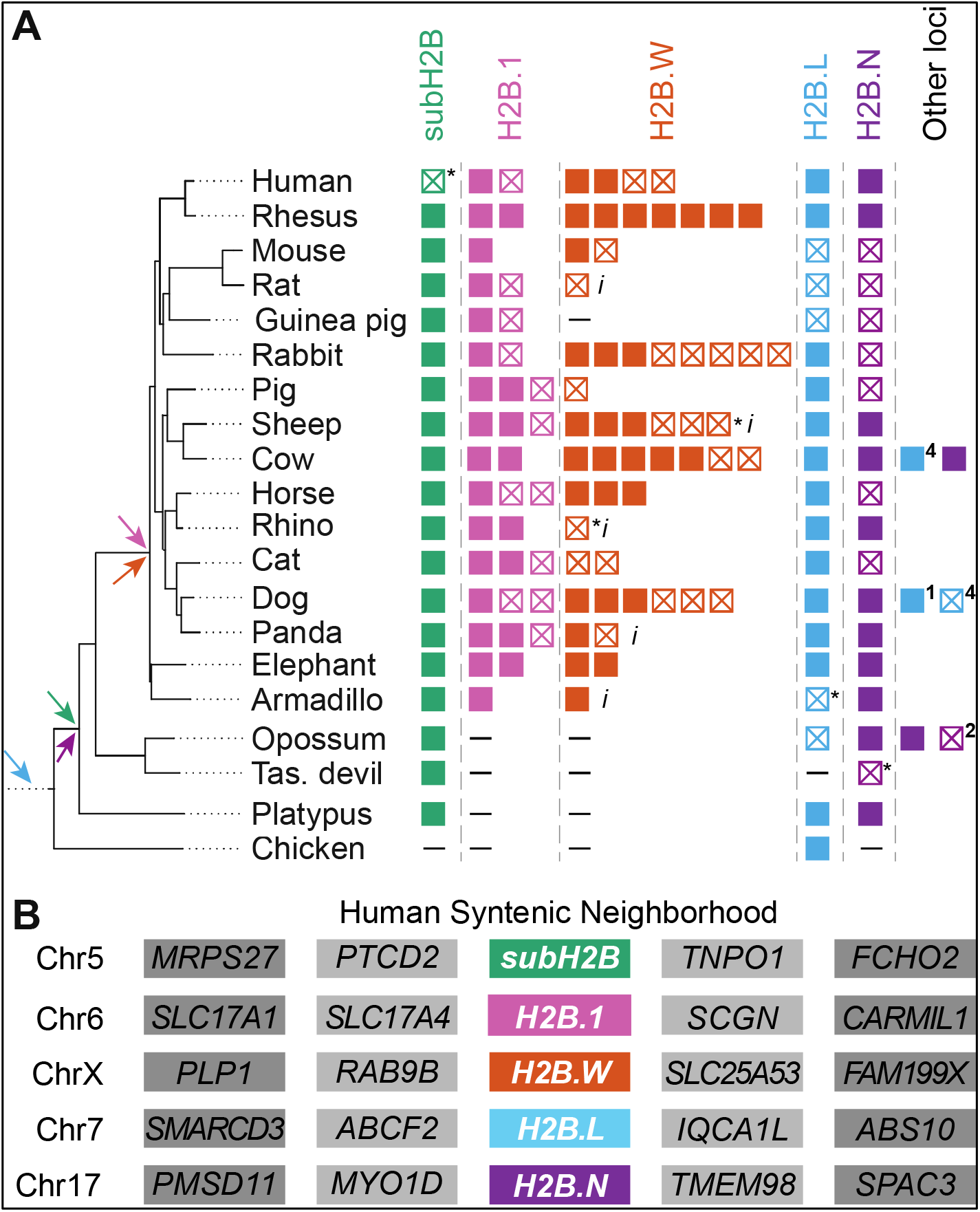
Retention and synteny of identified H2B variants in mammals. (**A**) A schematic representation of H2B variants and paralogs, along with a species tree of selected representative mammals and a non-mammalian outgroup, chicken, (Bininda-Emonds, et al. 2007). Colors distinguish H2B variants as in Figure 1A. Filled boxes represent intact open reading frames (ORFs) and empty boxes with a cross represent interrupted ORFs (inferred pseudogenes). An asterisk (*) indicates pseudogenization by a single nucleotide change which could either be sequencing error or a true mutation. A dash (–) indicates absence of histone within the syntenic location, and an “*i*” indicates incomplete sequence information. Colored arrows indicate the predicted origin of each H2B variant. Copies of variants found outside the syntenic neighborhood are shown as “other loci” with number of intact open reading frames and pseudogenes indicated. (**B**) A schematic of the shared syntenic genomic neighborhoods of each H2B variant in the human genome. All H2B variants in (A) were present in the same syntenic location across mammals (see Supplementary Figure S4 for detailed syntenic analyses) except “other loci”.

*H2B*.*L* is the only H2B variant for which we could identify an ortholog in the shared syntenic location in chicken, a non-mammalian outgroup (Figure 3A). We extended our analyses to other vertebrates and found *H2B*.*L* orthologs at least as far back as bony fishes in shared syntenic locations (Supplementary Figure S8). *H2B*.*L* orthologs from vertebrates also group with the previously identified ‘cleavage-stage dependent’ histones in sea urchin (Kemler and Busslinger 1986; Lai, et al. 1986; Marzluff, et al. 2006). Cleavage-stage histones, first described in sea urchin, are expressed at specific stages in embryogenesis. While orthologs of these sea urchin histones have been identified in other vertebrates (Ohsumi and Katagiri 1991; Mandl, et al. 1997; Tanaka, et al. 2001), our phylogeny lacks the bootstrap support for us to assign H2B.L and sea urchin cleavage-stage H2Bs as belonging to the same clade. The fragmented nature of the sea urchin genome assembly does not allow us to use shared synteny analysis to increase our confidence in assigning these to the same clade. Finally, the presence of an intron in all H2B.L orthologs but not in sea urchin cleavage-stage H2Bs challenges their orthology. Overall, our analyses suggests that four H2B variants arose in mammals, whereas *H2B*.*L* likely originated in the common ancestor of bony vertebrates (∼435 MYA), although this may even be an underestimate of its age.

### Rapid gene turnover of H2B orthologs

Next, we investigated the evolutionary dynamics of each H2B variant after its birth. We examined duplications, losses, and rates of protein sequence change. We found that all H2B variants, except *subH2B*, have experienced additional lineage-specific gene duplications (Figure 3A). *H2B*.*1* and *H2B*.*W* duplications occurred near or within the original syntenic locus in multiple mammals (Figure 3A, Supplementary Figure S7). In contrast, we found intronless duplicates of *H2B*.*L* and *H2B*.*N* in non-syntenic locations (other loci in Figure 3A), suggesting they arose via retrotransposition of the intron-containing progenitor genes. Notably, this pattern of gene duplication and retroposition often appears lineage-specific, with paralogs grouping with intron-bearing genes from the same species (Yang, et al. 2020). Based on this, we infer that *H2B*.*L* and *H2B*.*N* are likely to be expressed in the germline, since that is the only tissue in which retrogenes can be heritably integrated into the genome (Supplementary Figure S1).

Except for *H2B*.*1*, we found that no other H2B variant is universally retained in all mammals; they are all pseudogenized in at least one mammalian species (Figure 3A). For example, both *H2B*.*L* and *H2B*.*N* were pseudogenized in rodents. Our initial survey revealed that the human genome appears to encode a *subH2B* pseudogene, which is a single mutation away from encoding an intact open reading frame. Given the rarity of pseudogenization among mammalian *subH2B* genes, we investigated *subH2B* more closely across primates. We found that the frameshifting mutation (and subsequent early stop codon) found in humans is also present in chimpanzee, bonobo and gorilla suggesting that a true pseudogenization event occurred ∼9 million years ago in *Homininae* (Figure 3A, Figure 4, Supplementary Figure S9). We also found that *subH2B* pseudogenized at least five independent times in simian primates (Figure 4, Supplementary Figure S9). Thus, unusually among mammals, the subacrosomal subH2B variant appears to be non-functional in many primates.

**Figure 4.**
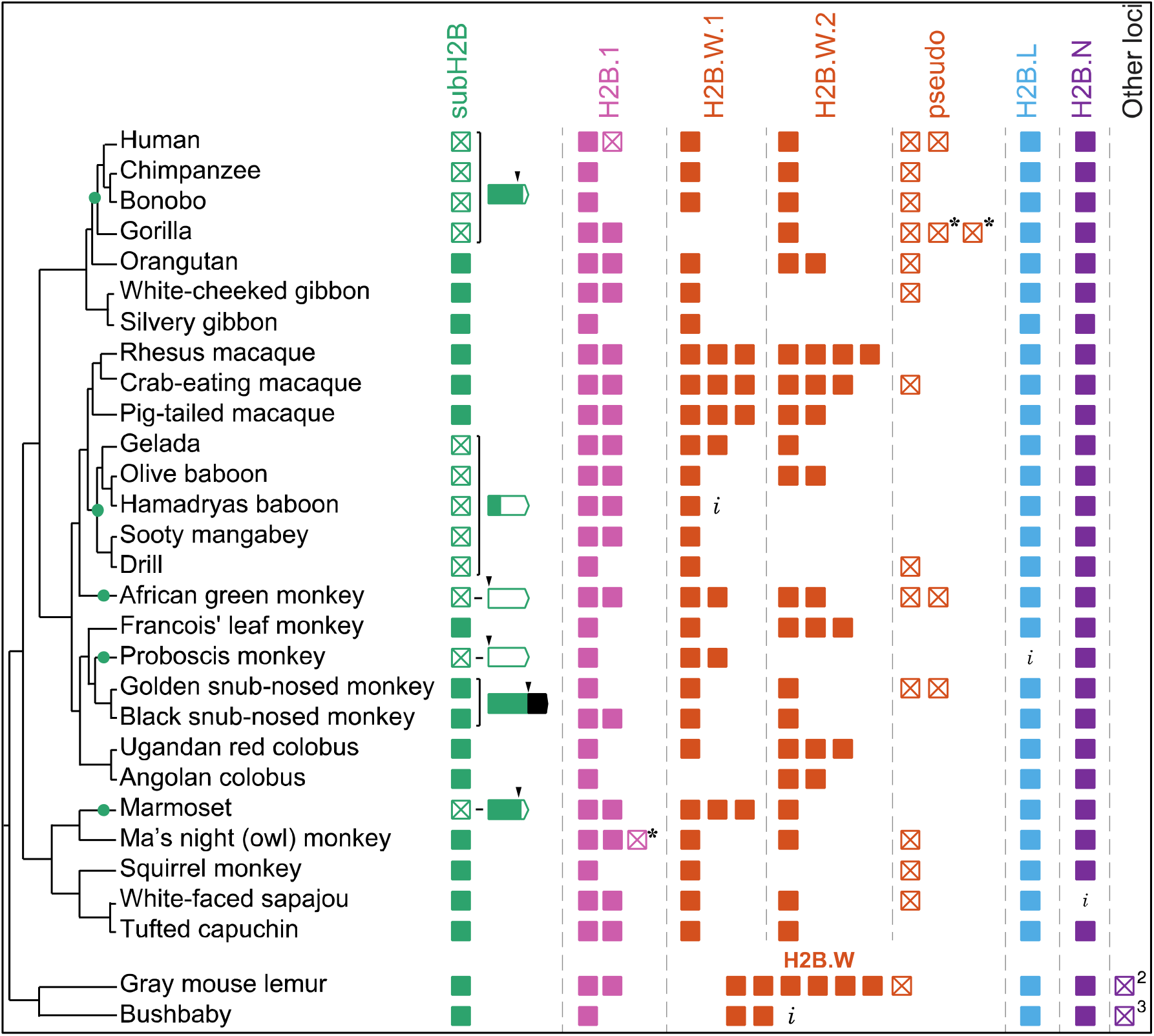
Evolutionary dynamics of H2B variants in primates. Schematic representation of identified H2B variants in primates, shown to the right of a species tree of primates (Perelman, et al. 2011; Osada 2015; Wright, et al. 2015). Colors distinguish H2B variants as in Figure 1A. Filled boxes represent intact open reading frames and empty boxes with a cross represent inferred pseudogenes. An asterisk (*) indicates pseudogenization by a single nucleotide change which could either be sequencing error or a true mutation. An “*i*” indicates incomplete sequence information. Copies of variants found outside the syntenic neighborhoods are shown as “other loci” with number of copies indicated. Green dots on the tree represent subH2B pseudogenization events inferred based on shared pseudogenizing mutations between species. For subH2B copies with pseudogenizing mutations or mutations that dramatically alter the ORF, disruptions to the ORF are schematized on a gene structure. Black arrowheads indicate nucleotide changes that result in a stop codon, frameshift or loss of start codon. Intact sequences in the ORF are filled green in the gene structure and disrupted sequences are empty. Black box indicates extension of ORF in snub-nosed monkeys. See Supplementary Figure S8 for more detailed description of subH2B mutations. H2B.W.1 and H2B.W.2 can be distinguished in simian primates using phylogeny (see Supplementary Figure S7) and so are shown in separate columns.

In contrast to *subH2B*, humans and other primates encode at least one intact copy of other H2B variants (Figure 4). Like most mammals, *H2B*.*L* and *H2B*.*N* are present in single copy in all primates, whereas *H2B*.*1* and *H2B*.*W* are present in multiple copies. Many primates have two copies of *H2B*.*1* that diverged from each other in the last common ancestor of simian primates (Supplementary Figure S10), although some species (including humans) subsequently lost one paralog. As we observed in our broader sample of mammals, *H2B*.*W* experienced dramatic duplications and pseudogenization in primates (Figure 4). However, unlike in other mammals, simian primate *H2B*.*W*.*1* and *H2B*.*W*.*2* genes can be readily distinguished by phylogenetic analyses of their HFDs. This suggests that primate *H2B*.*W*.*1* and *H2B*.*W*.*2* no longer experience gene conversion in their HFD and might have acquired partially non-redundant functions in primates. However, *H2B*.*W* gene turnover appears to be still active in primates; some primates have additional copies of H2B.W that do not reliably group with either H2B.W.1 or H2B.W.2, whereas other primates are missing an intact copy of either *H2B*.*W*.*1* or *H2B*.*W*.*2* (Supplementary Figure S11).

H2A and H2B form heterodimers before being incorporated into nucleosomes, suggesting that they might co-evolve. Previous work has identified dramatic diversification of H2A variants, especially short H2A variants in mammals (Govin, et al. 2007; Ferguson, et al. 2009; Shaytan, et al. 2015; Draizen, et al. 2016; Molaro, et al. 2018). However, with one exception, we did not observe any obvious correlations between the evolution of H2B and H2A variants when we examined their shared presence/absence in mammals. The one exception is that *H2A*.*1* and *H2B*.*1* are found in the same locus and share regulatory elements (Huh, et al. 1991). We found that a duplication of *H2B*.*1* is often accompanied by a duplication of *H2A*.*1* (Supplementary Figure S7). However, pseudogenization of one variant does not always lead to pseudogenization of the other. Thus, we cannot distinguish whether the apparent coevolution of *H2A*.*1* and *H2B*.*1* is due to genomic proximity and/or functional selection.

Overall, our phylogenomic studies of mammalian H2B variants reveal a dramatic, recurrent pattern of gene duplication and occasional functional loss. Lineage-specific loss of some H2B variants suggests that they are not essential for viability or fertility. Alternatively, the H2B variants might collectively perform an essential function but are individually functionally redundant.

### Evolutionary diversification and selective constraints acting on H2B variants

Given the long branch lengths of some H2B variants in our phylogeny (Supplementary Figure 1, 2) and the diversity revealed in some logo plots (Figure 2A), some H2B variants may have evolved more rapidly than RC H2B. To investigate this possibility, we compared the rate of protein divergence of RC H2B and H2B variants in a representative group of mammals spanning 100 million years of evolution (Figure 5A, Supplementary Table S2). For comparison purposes, we also included the *H2A*.*P* variant, which is one of the most rapidly diverging histone variants in mammals (Molaro, et al. 2018). We measured the pairwise identity of each mammalian H2B protein to its human ortholog (or orangutan ortholog for *subH2B*, since human *subH2B* is a pseudogene) and plotted it as a function of species divergence time (using TimeTree estimates, (Hedges, et al. 2015)). To be conservative, we chose the least divergent ortholog when multiple paralogs were found in the shared syntenic location. As expected, the highly conserved RC H2B shows the slowest rate of protein divergence. H2B.1 and H2B.L also evolve slowly. In contrast, H2B.N and subH2B exhibit an intermediate rate, whereas H2B.W shows the fastest rate of protein divergence among H2B variants, comparable to H2A.P (Figure 5A).

**Figure 5.**
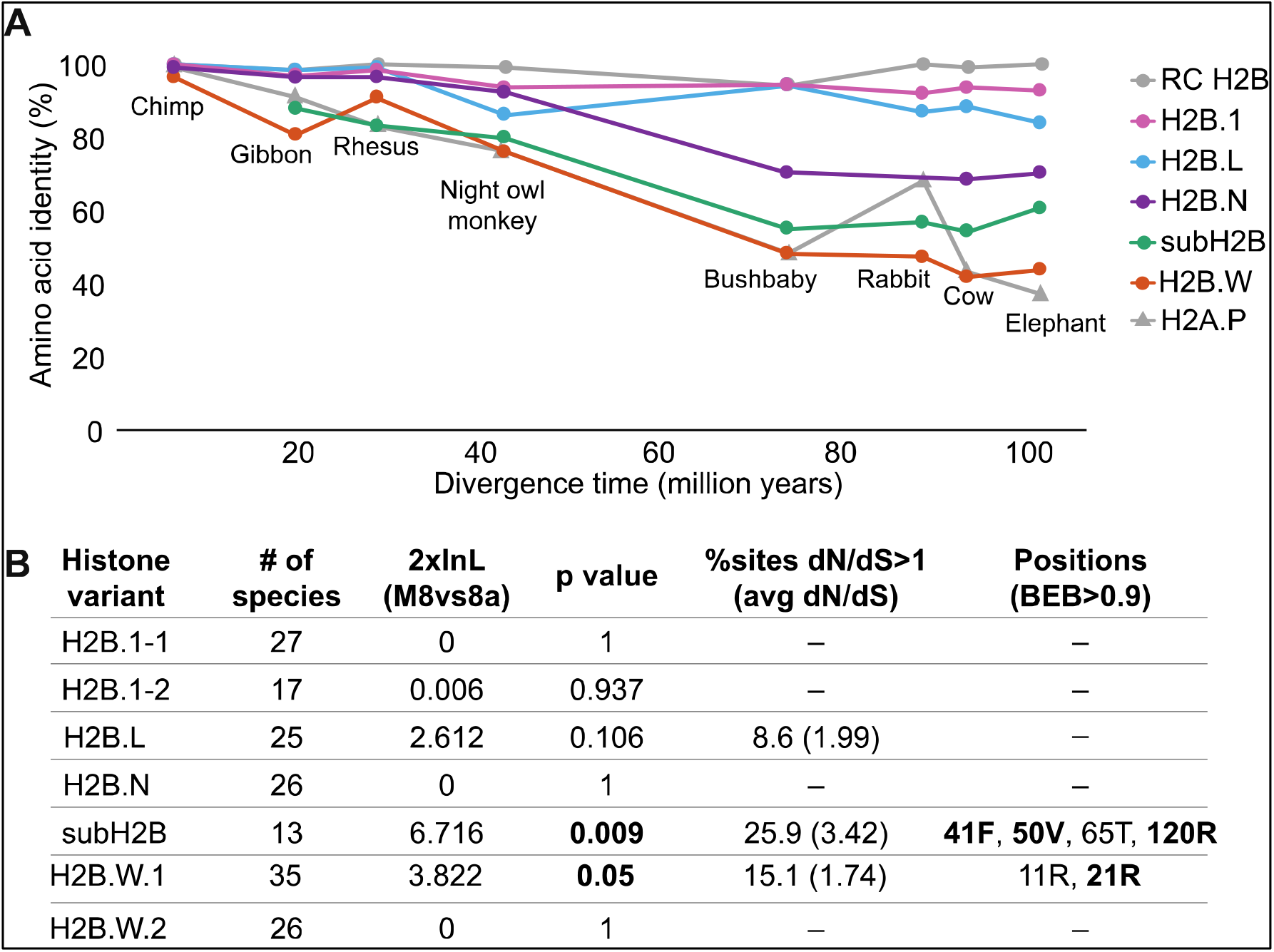
Evolutionary tempo of H2B variants in mammals. (**A**) Pairwise amino acid identity of H2Bs and H2A.P (Molaro, et al. 2018) showing comparisons of specified mammal orthologs versus either the human or orangutan (for subH2B) ortholog of each H2B variant. Percent identities (y-axis) are plotted against species divergence time (x-axis). For H2B variants with multiple copies, the copy with the highest identity to the human ortholog was used (See Supplementary Table S2 and Methods) to be conservative. (**B**) PAML analyses were used to look for site-specific positive selection (Supplementary Table S4). Log likelihood differences and p-values from the Model 8 vs Model 8a comparison are indicated. For the variants where likelihood tests suggest the presence of positive selection, the percentage of sites with *dN/dS*>1 is shown, along with the estimated average *dN/dS* for those sites. The ‘Positions’ column lists positively selected sites (M8 BEB>0.9); sites also identified by FUBAR analyses are highlighted in boldface (Murrell, et al. 2013). Amino acid residues shown correspond to the rhesus macaque protein sequence. See Supplementary Figure S12 for alignments showing positively selected residues.

The faster rates of amino acid change we observe could indicate diversifying (positive) selection at a select number of sites, or relaxed constraint. For example, complete lack of constraint would imply no functional selection for protein-coding capacity (*i*.*e*., neutrally evolving pseudogenes). To test for neutral evolution, we evaluated H2B variants by examining the ratio of rates of non-synonymous (amino acid altering, *dN*) to synonymous (*dS*) changes. Neutrally evolving sequences have *dN/dS* ratios close to 1, while strong purifying selection results in ratios near 0, with most non-synonymous changes disallowed. Using the same set of species selected above (Figure 5A) as input for PAML analysis (Yang 1997), we compared the relative likelihoods of models that assume sequences evolve neutrally versus those that allow non-neutral evolution. Specifically, we used PAML’s codeml program to estimate the likelihood of a simple evolutionary model (Model 0), where all sequences and all codons are assumed to have the same *dN/dS* ratio. We compared the likelihood of model 0 with *dN/dS* fixed at 1 (neutral) with that of model 0 with *dN/dS* estimated from the alignment. Using this test, we found strong evidence for purifying selection for all H2B variants across mammals (Supplementary Data Table S3), rejecting neutrality and the possibility that their high protein divergence is due to pervasive pseudogenization. Furthermore, the overall *dN/dS* estimates from these analyses are consistent with the relative evolutionary rates of the H2B variants based on protein divergence (Figure 5A).

The signatures of overall purifying selection in the H2B variants do not rule out the possibility that a subset of sites might nevertheless evolve under positive selection (*dN/dS* >1). Indeed, H2B variants in plants (Jiang, et al. 2020) and short H2A variants in mammals (Molaro, et al. 2018) show evidence of both overall purifying selection and positive selection at selected sites. To investigate this possibility, we analysed H2B variant sequences from simian primates, a clade with a level of evolutionary divergence that is ideal for codon-by-codon analyses. We analysed intact ORF sequences from 27 species for most variants. However, we had only 13 sequences for subH2B since it is pseudogenized in many primates (Figure 4, Supplementary Figure S9). We performed maximum likelihood analyses using PAML (Yang 1997) and FUBAR (HyPhy package (Pond, et al. 2005; Murrell, et al. 2015)) to investigate whether a subset of codons experience positive selection. Using PAML, we identified signatures of positive selection for *subH2B* and *H2B*.*W*, but not for the other variants. We found a strong signature of diversifying selection for *subH2B*, with an estimated 25.9% of sites evolving with an average *dN/dS* of 3.42; four sites in the HFD showed high posterior probabilities of evolving under positive selection (Figure 5B, Supplementary Table S4, Supplementary Figure S12). An analysis of 35 *H2B*.*W*.*1* paralogs/orthologs from simian primates also revealed diversifying selection, with an estimated 15.1% of sites evolving with an average *dN/dS* of 1.74 (Figure 4B, Supplementary Table S4). However, only one site in the H2B.W.1 N-terminal tail showed a high posterior probability of positive selection (Figure 5B, Supplementary Figure S12). FUBAR analyses also identified additional sites in subH2B and H2B.W that might have undergone positive selection (posterior probability>0.9) (Supplementary Table S4). Unlike PAML analyses, FUBAR analyses identified *H2B*.*W*.*2, H2B*.*1, H2B*.*L* and *H2B*.*N* as also having undergone diversifying selection in simian primates, along with *subH2B* and *H2B*.*W* (Supplementary Table S4). Our limited understanding of functional residues in H2B variants prevents us from making any informative biological predictions about the rapidly evolving sites. Overall, our findings of strong purifying selection suggests that H2B variants perform vital functions, whereas our findings of positive selection suggest that they have been subject to recurrent genetic innovation.

### H2B variants are expressed in mammalian germlines

Most H2B variants in this study remain functionally uncharacterized. To begin to explore their function, we examined their expression in mammals. Similar analyses had previously revealed putative germline-specific functions of many rapidly evolving H2A variants (Molaro, et al. 2018; Molaro, et al. 2020). Prior studies have shown that some H2B variants are primarily expressed in testes of rodents, bull, or human. For others, including the novel variants identified in this study, the site of expression is not known.

We examined publicly available RNA-seq data for the expression of all H2B variants in diverse somatic (brain, liver, kidney, heart) and germline (testes and ovaries) tissues from a wide range of species – opossum, dog, pig, mouse, human and chicken (Supplementary Table S5, Methods). For comparison, we included a previously published “housekeeping” gene (*C1orf43*) that is ubiquitously expressed (Eisenberg and Levanon 2013). We did not detect expression of any H2B variants in somatic tissues (Supplementary Figure S13A-E) but did detect robust expression in the majority of germline samples.

The subH2B variant protein was originally isolated from bull and rodent sperm (Aul and Oko 2001). In our RNA-seq analysis, *subH2B* is abundantly expressed in testes of representative mammals, except humans, where it is pseudogenized (Supplementary Figure S13A-G). Although the pseudogenizing mutation in humans occurred relatively recently (Figure 4), absence of detectable *subH2B* implies either that its regulatory sequences are also non-functional or that its transcript is rapidly degraded in human cells. In contrast, we observed low levels of *subH2B* transcript in the testes of rhesus macaque, where the ORF is intact (Supplementary Figure S13H).

Consistent with previous work in mice (Branson, et al. 1975; Shires, et al. 1975; Zalensky, et al. 2002; Govin, et al. 2007; Montellier, et al. 2013), we found that *H2B*.*1* is expressed in testes and ovaries of all mammals we analysed (Supplementary Figure S13B-G). However, in contrast to previous studies that reported H2B.W.1 expression in human sperm (Churikov, et al. 2004; Boulard, et al. 2006), we detected no or very low levels of *H2B*.*W* expression in all species we examined (Supplementary Figure S13B-G). We note that expression of human H2B.W.1 and H2B.W.2 protein is enriched in sperm samples in a publicly available expression database (Human Protein Atlas) (Uhlén, et al. 2015; Thul, et al. 2017; Uhlen, et al. 2017) further supporting their expression in the male germline. Finally, we found that both *H2B*.*N* and *H2B*.*L* are expressed in ovaries of opossum, dog, and humans, whereas *H2B*.*L* is expressed in both testes and ovaries in pigs (Supplementary Figure S13A-G). We did not examine the expression of either *H2B*.*N* and *H2B*.*L* in mice, or of *H2B*.*N* in pigs, because these genes have multiple pseudogenizing mutations in these species (Supplementary Figure S13C, D). We also found *H2B*.*L* expression in chicken ovaries (Supplementary Figure S13I), demonstrating that ovarian expression of *H2B*.*L* likely predates the divergence of birds and mammals.

We were concerned about the inconsistency between our and previous analyses about the expression of some H2B variants (especially *H2B*.*W*). We speculated that this inconsistency might be due to unusual RNA structures or tissue heterogeneity. Instead of poly-A tails, RC histone transcripts have unusual stem-loop RNA structures at their 3’ ends that bind stem-loop binding protein (SLBP), which regulates their stability and translation (Dávila López and Samuelsson 2008; Marzluff, et al. 2008). Because of this, RC histones are typically under-represented in poly(A)-selected RNA-seq datasets. In contrast to RC histones, most histone variants are thought to have polyadenylated transcripts and lack stem loops. Yet, previous work has suggested that RC histones can have alternate mRNA processing modes (Molden, et al. 2015). To investigate this dichotomy in RNA structure further, we searched for stem loop structures and poly(A) signals close to the stop codons of all H2B variant genes. Stem loop sequences are easily recognized, whereas poly(A) signal detection is less accurate, with false positive and false negative findings. We were able to detect a poly(A) signal in the 3’ UTR of most *subH2B, H2B*.*N* and *H2B*.*L* genes (Supplementary Figure S14). Unexpectedly, we detected both stem-loop and poly(A) sequences in the 3’ UTRs of the *H2B*.*1* and *H2B*.*W* genes (Supplementary Figure S14). Our analyses further reveal that histone variants may also be subject to alternate processing; this layer of histone processing and regulation has been poorly studied. We speculate that alternate RNA processing may operate for some H2B variant genes, and that the presence of stem loops might have affected our ability to detect *H2B*.*W*.*1* and *H2B*.*W*.*2* transcripts in publicly available RNA-seq datasets that mostly use poly(A) selection.

A second challenge for detecting histone variant expression in RNA-seq analyses could be cell heterogeneity. For example, many different cell types and developmental stages are present in testes and ovaries. Bulk RNA-seq analyses may be unable to detect robust expression if H2B variants are only transcribed in a small subset of cells. To more closely investigate this possibility, we examined expression of H2B variants during human spermatogenesis. We detected robust expression of *H2B*.*1* in sperm, with expression increasing during early stages of spermatogenesis (Figure 6A, Supplementary Figure S13F) but decreasing post-meiosis, consistent with previous reports (van Roijen, et al. 1998; Govin, et al. 2007; Montellier, et al. 2013). In contrast, we did not detect expression of *H2B*.*W*.*1* or *H2B*.*W*.*2* in either spermatogenesis or oogenesis datasets (Figure 6A, B, Supplementary Figure S13F, G). It is possible that both *H2B*.*W* variants are expressed at stages of gametogenesis (Churikov, et al. 2004) that are not captured in our data analyses due to lack of poly(A) tails at the 3’ end of their transcripts (above). Alternatively, even low expression of *H2B*.*W* variants may be sufficient for their function in sperm.

**Figure 6.**
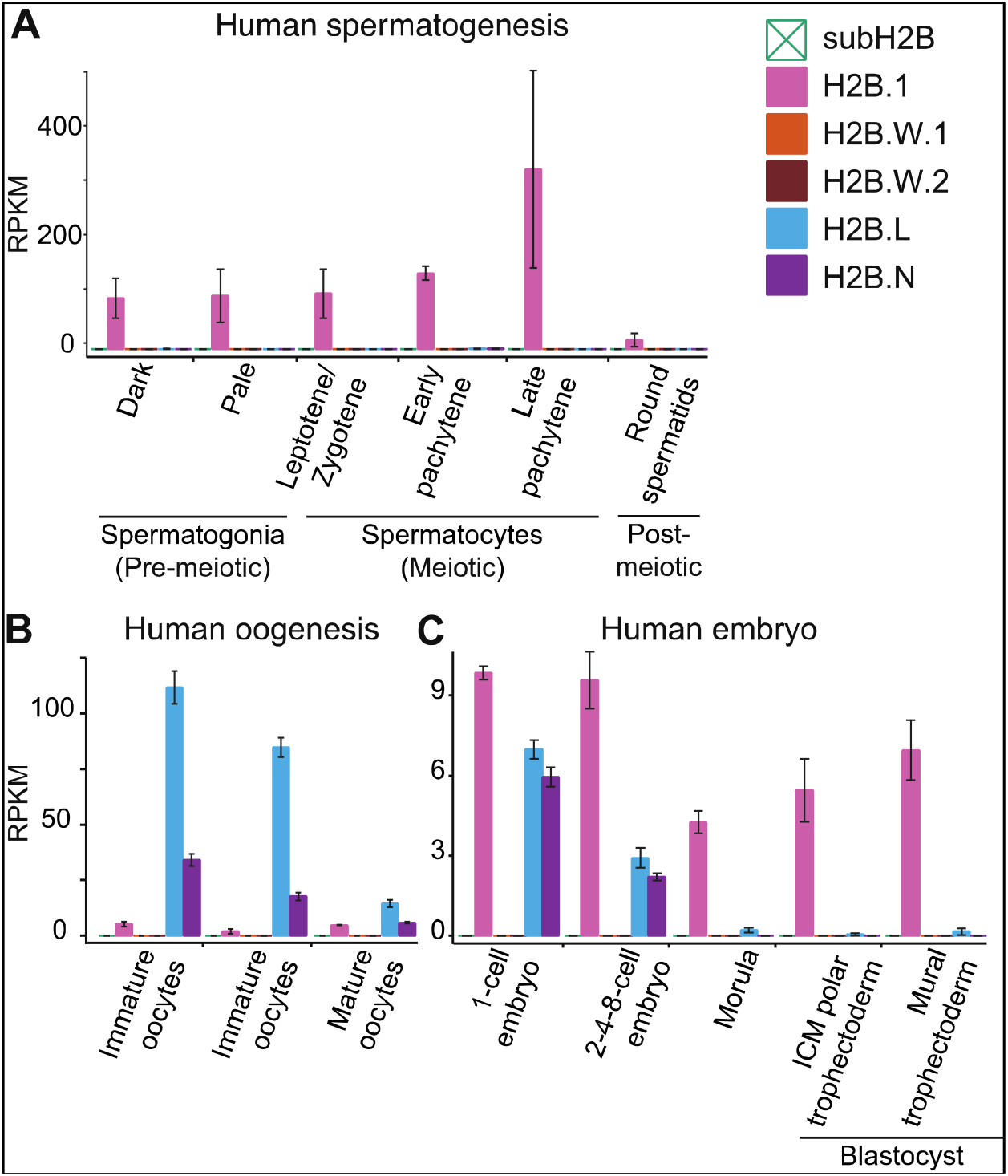
H2B variant expression during human gametogenesis and embryogenesis. RNA expression of H2B variants (reads per kilobase per million mapped reads, RPKM) in publicly available bulk RNA-seq data across different stages of (**A**) human spermatogenesis, (**B**) human oogenesis and (**C**) human embryogenesis. Legend shows colors corresponding to each histone variant (note that subH2B is pseudogenized in humans). The bar heights show median RPKMs of biological replicates and error bars show median absolute deviations. See Supplementary Figure S10 for additional analyses of H2B variants’ expression in somatic and reproductive tissues of other mammals.

Analyses of human oogenesis revealed robust expression of *H2B*.*L* and *H2B*.*N* in oocytes, with levels increasing across oogenesis (Figure 6A, B, Supplementary Figure S13F, G). Neither *H2B*.*L* nor *H2B*.*N* were detected in granulosa cells, which are the somatic cells of the female germline, suggesting again that expression is restricted to the germline. We also detected low expression of *H2B*.*1* in human oogenesis (Figure 6B), consistent with previous analyses of mouse oogenesis (Montellier, et al. 2013; Beedle, et al. 2019). Finally, we detected expression of *H2B*.*1* (consistent with a previous study in mice (Montellier, et al. 2013)), *H2B*.*L*, and *H2B*.*N*, but not of any of the other H2B variants during embryogenesis (Figure 6C, Supplementary Figure S13G). Overall, our analyses suggest that expression of most H2B variants is restricted to the male germline. However, newly identified histones *H2B*.*N* and *H2B*.*L* are primarily expressed in ovaries and early embryos, where they may play key roles in female fertility and early development like the cleavage stage histones of sea urchins (Poccia, et al. 1981; Tanaka, et al. 2001; Oliver, et al. 2003).

## Discussion

Histones perform the critical task of packaging genomes and regulating important DNA-based processes (*e*.*g*., transcription, DNA repair, chromosome segregation) in most eukaryotes. Because of their critical genome-wide functions, many replication-coupled (RC) histones evolve under extreme evolutionary constraint, permitting only limited changes in protein sequence even over nearly a billion years of divergence between protists and humans. In contrast, histone variants can acquire changes that enable them to elaborate new functions, contributing to the eukaryotic nucleosome’s remarkable structural and functional plasticity. Understanding the evolutionary history of histones can reveal how biological challenges that different organisms face have been resolved by chromatin innovation.

We reveal an extensive repertoire of H2B variants in mammalian lineages, including three previously undescribed histone variants (H2B.O, H2B.L, H2B.N). Two H2B variants are only found in a small subset of mammals – H2B.E in rodents and H2B.O in platypus – whereas five variants (subH2B, H2B.1, H2B.N, H2B.L, H2B.W) are found more extensively across eutherian mammals. We find that one of the newly discovered variants, H2B.L, arose prior to the origin of bony vertebrates. Given its age and widespread retention, it is somewhat surprising that H2B.L has escaped detection until now. We attribute this to the difficulty of correctly classifying histone variants when interrogating single genomes (*e*.*g*., human or mouse). The atypical absence of H2B.L (and H2B.N) from the mouse genome, where the most extensive characterization of variant histones has been carried out, likely exacerbated this difficulty (Aul and Oko 2001; Govin, et al. 2007; Montellier, et al. 2013; Shinagawa, et al. 2015). Both of these reasons further highlight the value of comprehensive phylogenomic studies in identifying and classifying meaningful functional innovation in histone variant genes.

H2B variants display a range of evolutionary divergence rates across mammals. H2B.1 and H2B.L evolve slowly, whereas subH2B, H2B.N and H2B.W evolve more rapidly. We also detected signatures of positive selection for a subset of residues in several H2B variants in simian primates. These observations are consistent with previous work that suggests male germline-specific genes tend to evolve more rapidly (Retief and Dixon 1993; Wyckoff, et al. 2000; Torgerson, et al. 2002; Turner, et al. 2008; Martin-Coello, et al. 2009). In addition to divergence in their HFD, H2B variants show significant divergence from RC H2B in their N- and C-terminal tails, so much so that many H2B variant tails cannot be reliably aligned with RC H2B. The most dramatic changes were seen for H2B.W variants, which have significantly longer N and C-terminal tails, and H2B.N variants, which have much shorter C-terminal tails; these changes could significantly impact nucleosome packaging and stability. Furthermore, these differences in the tail could contribute to altered PTMs on the tails that are crucial to chromatin interactions and their regulation. Even the relatively conserved H2B.1 variant differs from RC H2B at some sites that can be post-translationally modified (Zalensky, et al. 2002; Li, et al. 2005; Lu, et al. 2009) to facilitate looser packaging of chromatin than by RC H2B (Rao and Rao 1987; Singleton, et al. 2007; Urahama, et al. 2014). Thus, the newly identified H2B variants represent a rich source of additional structural, functional, and regulatory complexity.

All variants except H2B.1 have been lost in at least one mammalian genome. This suggests that all H2B variants except H2B.1 might be dispensable for viability and/or fertility. Moreover, even loss of H2B.1 in knockout mice can be compensated for by RC H2B with unusual PTMs that allow for similar nucleosome structure as H2B.1 (Montellier, et al. 2013). Yet, double mutant mice of H2B.1 and H2A.1 are infertile and inviable (Shinagawa, et al. 2015). One explanation for this inviability is that while RC H2B can compensate for H2B.1 function, RC H2A does not appear to compensate for H2A.1, leading to a stoichiometric imbalance (Shinagawa, et al. 2015). Since the properties of variant nucleosomes can be affected by variation in any of the four histone components, this represents an additional layer of chromatin complexity and innovation that remains almost entirely unexplored.

Whereas H2B.E is expressed in neurons (Santoro and Dulac 2012), all other mammalian H2B variants have germline-biased expression. SubH2B, H2B.1, H2B.W, and H2B.O are expressed in testis/sperm while H2B.1, H2B.N and H2B.L are expressed in ovaries/oocytes and embryos. Together with the invention of germline-enriched short H2A variants in mammals (Molaro, et al. 2018), our study reiterates the important role of evolutionary innovation of chromatin functions in mammalian germ cells, similar to what has been previously observed in plants (Jiang, et al. 2020). Spermatogenesis may require constant chromatin innovation since it is a hotbed of genetic conflicts (Moore and Haig 1991; Moore and Reik 1996; Torgerson, et al. 2002; Civetta and Ranz 2019), both within genomes (*e*.*g*., transposable elements, post-meiotic segregation distortion) and between genomes (*e*.*g*., sperm competition during fertilization). Given its subacrosomal localization, subH2B is more likely to play a role in gamete fusion rather than in chromatin. However, subH2B appears to be non-functional in most simian primates and its function remains uncharacterized in any mammal.

Spermatogenesis in mammals and many other animal species also involves the near-complete replacement of histones by highly-charged basic proteins, called protamines, which ensure tight DNA packaging in sperm heads (Oliva and Dixon 1991; Hammoud, et al. 2009). Following fertilization, the paternal genome must be stripped of protamines and repackaged by histones, such that paternal and maternal genomes (which never undergo protamine replacement) can initiate embryonic cell cycles in an orderly fashion. Deposition of H2B.1 is a key transitional step between RC histone and protamine-packaged genomes in early spermatocytes (Montellier, et al. 2013; Shinagawa, et al. 2015) and for repackaging the protamine-rich paternal genome into histones following fertilization (Montellier, et al. 2013). Notably, even after the protamine transition, 10-15% of basic nuclear proteins in mature sperm still constitute histones (Tanphaichitr, et al. 1978; Rousseaux, et al. 2005). Even though H2B.1 is removed from mature sperm, other functionally uncharacterized H2B variants may be required for as-yet-unknown critical functions for spermatogenesis. For example, ectopically expressed H2B.W.1 appears to localize to telomeric chromatin in cell lines (Churikov, et al. 2004), suggesting it might play a ‘bookmarking’ role, as has been hypothesized for histones post-fertilization (Hammoud, et al. 2009). Although several H2B variants appear to be highly enriched in the male germline, this does not eliminate the possibility that they might play important roles during oogenesis and early embryogenesis. For example, the H2A.B variant is testis-enriched in expression but nevertheless plays key roles in oogenesis and post-implantation development in mice (Molaro, et al. 2020).

In contrast to spermatogenesis, oogenesis does not appear involve dramatic chromatin changes analogous to the protamine transition. Yet maternal inheritance of certain histone variants is essential for embryonic viability and development (Martire and Banaszynski 2020). During oogenesis, chromatin undergoes chromosome condensation (Bogolyubova and Bogolyubov 2020), withstands double-stranded breaks during meiotic recombination, and survives a long meiotic arrest in mammals (Cheng, et al. 2009; Lake and Hawley 2012; Carroll and Marangos 2013). In addition, histone variants may “mark” imprinted regions of the inherited maternal genome in embryos. Reprogramming of maternal genomes to match the epigenetic state of paternal genomes is also critical for fitness in many animals (Potok, et al. 2013). Maternally deposited histone variants may also be crucial for the initial stages of embryogenesis, especially to mediate the protamine-to-histone transition of the paternal genome and post-translational modifications of histones for zygotic genome activation. Despite these specialized chromatin requirements, very few chromatin innovations have been described for mammalian oogenesis, unlike for spermatogenesis. So far, only a few oocyte-specific variants, including some linker histone H1 variants, have been described (Martire and Banaszynski 2020; Talbert and Henikoff 2021). Although some H2A variants, including H2A.1, H2A.B and macroH2A, are expressed during oogenesis, their functions, and their interactions with H2B variants remain uncharacterized. Our identification of two previously uncharacterized female germline-enriched histone variants — H2B.L and H2B.N — could thus reveal important insights into chromatin innovation and requirements during oogenesis. The oogenesis-expressed H2B variants (*H2B*.*1, H2B*.*L*, and *H2B*.*N*) are also detected in human embryos, suggesting the possibility of embryonic functions, which could be elucidated by future *in vivo* analyses.

The newly identified H2B variants also present some novel features that have not been previously observed in histones. Although H2B.L resembles RC H2B in its HFD, its highly diverged N-terminal tail includes a polyglutamine repeat and overall lower charge, suggesting it may confer different functionality and looser chromatin packing when incorporated into nucleosomes. Its near-ubiquitous presence across vertebrate genomes and strong sequence conservation motivates future functional studies. In contrast to H2B.L, H2B.N is dramatically different from RC H2B. For example, most H2B.N proteins have a significant C-terminal truncation that removes the αC domain, eliminating the important nucleosome acidic patch that mediates many other chromatin interactions (McGinty and Tan 2021). This feature is so unusual that that it raises the possibility of a non-nucleosomal function for H2B.N (like subH2B), which could be revealed by biochemical and cytological analyses.

Our analyses not only reveal chromatin innovation in mammalian germlines, but may also provide important clues for chromatin aberrations that can arise in cancer cells. For example, misexpression of other germline-specific histone variants and mutations in RC H2B can be detected in cancer cells (Bennett, et al. 2019; Nacev, et al. 2019; Bagert, et al. 2021; Chew, et al. 2021). Recent studies have identified H2B.W.2 as a potential driver gene in cervical cancer (Xu, et al. 2021). Our phylogenomic analyses thus pave the way for future functional studies of H2B variants in gametogenesis and their misexpression in somatic cells, with implications for cancer and other diseases.

## Methods

### Identification of H2B variants

To identify mammalian H2B variants we iteratively queried the assembled genomes of eighteen mammals– human (*Homo sapiens*), mouse (*Mus musculus*), rat (*Rattus norvegicus*), guinea pig (*Cavia porcellus*), rabbit (*Oryctolagus cuniculus*), pig (*Sus scrofa*), sheep (*Ovis aries*), cow (*Bos taurus*), horse (*Equus caballus*), white rhinoceros (*Ceratotherium simum*), cat (*Felis catus*), dog (*Canis lupus familiaris*), panda (*Ailuropoda melanoleuca*), elephant (*Loxodonta africana*), armadillo (*Dasypus novemcinctus*), opossum (*Monodelphis domestica*), Tasmanian devil (*Sarcophilus harrisii*), and platypus (*Ornithorhynchus anatinus*), as well as a non-mammalian outgroup species, chicken (*Gallus gallus*) (Supplementary Table S1). We used TBLASTN (Altschul, et al. 1990; Altschul, et al. 1997) on each species’ genome to perform a homology-based search starting with human H2B.W.1 (Q7Z2G1) (Supplementary Table S1) as our query. We chose H2B.W.1 as a query sequence instead of an RC H2B to focus our search on more divergent H2B genes and because most mammalian genomes encode many near-identical RC H2B sequences. To ensure that we had not missed any divergent H2B homologs, we repeated our analyses using all H2B variants in this study as queries in TBLASTN searches and did not retrieve additional hits. To determine the age of H2B.L, we searched for H2B.L via TBLASTN in non-mammalian vertebrates– zebra finch (*Taeniopygia guttata*), western clawed frog (*Xenopus tropicalis*), coelacanth (*Latimeria chalumnae*), zebrafish (*Danio rerio*), elephant shark (*Callorhinchus milii*), lamprey (*Petromyzon marinus*), sea urchin (*Strongylocentrotus purpuratus*), and fruit fly (*Drosophila melanogaster*) (Supplementary Table S1). H2B variant orthologs in 29 primates were identified using TBLASTN analyses of NCBI’s non-redundant nucleotide collection (nr/nt) and whole-genome shotgun contigs (wgs) databases (Supplementary Table S1).

Once histone variants were identified, we used shared synteny (conserved genetic neighborhood) to identify putative orthologs in all representative mammalian genomes. We retrieved nucleotide sequences for all hits and their genomic neighborhoods, and recorded coordinates for syntenic analyses using the UCSC Genome Browser (Kent, et al. 2002). For syntenic analyses (Figure 3B, Supplementary Figures S3, S7, S8), 2-3 annotated genes on either side of each histone variant were identified (flanking genes) from mouse or human genomes. TBLASTN searches using each flanking gene were performed to identify orthologs and therefore the syntenic regions in all selected mammalian genomes and chicken. In some genomes, the syntenic location was split between multiple scaffolds (double slashes in figures). *H2B*.*L* could not be identified in Tasmanian devil, likely because the syntenic region is split between two scaffolds: the intervening sequence may be missing from the assembly. Histones or flanking genes located on scaffolds labeled with Chr_UN were not included in our analyses.

Since RC H2Bs are present in numerous identical copies in mammalian genomes, we only used one copy of any identical RC H2Bs in our analyses. We used a copy of RC H2B that is present in six copies in the human genome (H2Bc4/ H2Bc6/ H2Bc7/ H2Bc8/ H2Bc10) and three copies in the mouse genome. Exons and introns in variants *H2B*.*W, H2B*.*L* and *H2B*.*N* were annotated based either on protein alignments with closely-related species or on Ensembl, RefSeq or GenScan predictions. Since the N-and C-terminal residues of H2B.W orthologs show high divergence in mammals, our current annotations in non-primate species may need to be revised with further experimental evidence. Pseudogenes were annotated based on disrupted open reading frames or the presence of gene remnants as determined by a TBLASTN search of the histone variant sequence against its syntenic regions (as in the case of *H2B*.*W* and *H2B*.*1*). In three cases, H2B variant copies were also found on the same chromosome immediately outside the syntenic location– one cow and sheep *H2B*.*W* variant and a horse *H2B*.*1* pseudogene. These are not annotated as other loci in Figure 2A since they were found on the same chromosome as the ancestral gene near the syntenic region.

We used shared synteny, sequence similarity and phylogenetic analyses (below) to classify ORFs and pseudogenes into H2B variant families (H2B.E, subH2B, H2B.1, H2B.W, H2B.L, and H2B.N).

### Phylogenetic analyses

All protein and nucleotide alignments were performed using the MUSCLE algorithm (Edgar 2004) in Geneious Prime 2019.2.3 (https://www.geneious.com) and all phylogenies were generated using maximum-likelihood methods in PhyML (Guindon and Gascuel 2003; Guindon, et al. 2010) with 100 bootstrap replicates. Since RC H2B are present in many near-identical copies in mammalian genomes, we used a random number generator to select two arbitrary copies of H2B from each species. Our protein phylogenies used alignments of either the HFD and αC domain, or the full-length sequences, with the Jones-Taylor-Thornton substitution model (Jones, et al. 1992). Our nucleotide phylogenies used the HKY85 substitution model (Supplementary Figure S10, S11). Pseudogenes were not included in any tree.

Sequences of *H2B*.*W* from mammals were analysed for evidence of recombination using the GARD algorithm at datamonkey.org (Kosakovsky Pond, et al. 2006).

### Calculating rate of protein divergence for histones

We used full-length protein sequences of all H2B variants to calculate pairwise identities between representative mammal orthologs (Figure 5A; Supplementary Table S2). We used the human ortholog as a reference for all H2B variants, except subH2B, which has been pseudogenized in humans; therefore, we used orangutan subH2B as a reference sequence. Sequence divergence levels for H2A.P were obtained from a previous study (Molaro et al., 2018). We obtained median species divergence times from the TimeTree database (www.timetree.org) (Hedges et al. 2015).

### Analysis of evolutionary selective pressures

We analyzed selective pressures on H2B variants in diverse mammals or in simian primates using the codeml algorithm from the PAML suite (Yang 1997) (Supplementary Table S3, S4). For all tests, we generated codon alignments using MUSCLE (Edgar 2004), and manually adjusted them to improve alignments if needed. We also trimmed sequences to remove alignment gaps and segments of the sequence that were unique to only one species. We found no evidence of recombination for any of these alignments using the GARD algorithm at datamonkey.org (Kosakovsky Pond, et al. 2006). We used the alignment to generate a tree using PhyML maximum-likelihood methods with the HKY85 substitution model (Guindon et al. 2010).

To test for gene-wide purifying selection (Supplementary Table S3), we used codeml’s model 0, which assumes a single evolutionary rate for all lineages represented in the alignment. We compared likelihoods between model 0 with a fixed *dN/dS* value of 1 (neutral evolution) and model 0 with *dN/dS* estimated from the alignment. We determined statistical significance by comparing twice the difference in log-likelihoods between the two models to a χ^2^ distribution with 1 degree of freedom (Yang 1997).

To test whether a subset of residues evolves under positive selection (Supplementary Table S4), we compared nested pairs of “NSsites” evolutionary models. We compared likelihoods between NSsites model 8 (where there are 10 classes of codons with dN/dS between 0 and 1, and an eleventh class with dN/dS >1) and either model 7 (which disallows dN/dS to be equal to or exceed 1) or model 8a (where the eleventh class has dN/dS fixed at 1). We determined statistically significance by comparing twice the difference in log-likelihoods between the models (M7 vs M8 or M8 vs M8a) to a χ^2^ distribution with the degrees of freedom reflecting the difference in number of parameters between the models being compared (Yang 1997). For alignments that showed statistically significant support for a subset of sites under positive selection, sites with a Bayes Empirical Bayes posterior probability >90% in M8 were classified as positively selected sites.

In addition to PAML, we used the FUBAR (Murrell, et al. 2013) and BUSTED (Murrell, et al. 2015) algorithms from datamonkey.org to estimate selection at each site or on the whole gene respectively (Supplementary Table S4).

### Logo plots and nucleosome structure

Logo plots were generated using WebLogo (weblogo.berkeley.edu; (Crooks, et al. 2004)) using one copy of each H2B variant or RC H2B protein sequences from each of the following species: sheep, dog, elephant, cow, bushbaby (*Otolemur garnettii*), mouse lemur (*Microcebus murinus*) and rhesus macaque (*Macaca mulatta*). We calculated a two-way Jensen-Shannon distance metric (Doud, et al. 2015) at each amino acid position in the HFD and the αC domain as a quantitative estimate of conservation of each residue between each H2B variant and RC H2B. We also compared of all H2B variants together versus RC H2B as previously described (Molaro, et al. 2018) to identify residues that differ between RC H2B and all H2B variants. This analysis did not reveal any residues common to all H2B variants and distinct from RC H2B.

We used Phyre2 (Kelley, et al. 2015) to construct a homology model of the HFD of H2B variants. This software used existing H2B crystal structures to model the structure of human H2B variant protein sequences (or rhesus macaque for subH2B). We used the Chimera software (Pettersen, et al. 2004) to display the resultant predicted models with high confidence and highlighted residues of interest on a previously published human nucleosome structure (PDB:5y0c) (Arimura, et al. 2018). The isoelectric point and charge for human H2B variants (Supplementary Figure S6C) was computed using Protpi (https://www.protpi.ch/).

### RNA-seq analysis

We analyzed publicly available transcriptome data from chicken, opossum, dog, pig, mouse and human to approximately quantify expression of H2B variants in somatic and germline tissues (Supplementary Table S5). We downloaded FASTQ files using NCBI’s SRA toolkit (https://www.ncbi.nlm.nih.gov/books/NBK158900), and mapped reads to same-species genome assemblies using the STAR mapper (Dobin, et al. 2013). We used the “-- outMultimapperOrder Random --outSAMmultNmax 1 --twopassMode Basic” options so that multiply mapping reads were assigned randomly to a single location. We then used genomic coordinates of each ORF and the BEDTools multicov tool (Quinlan and Hall 2010) to count reads overlapping each gene. We then used R (R Core Team 2015) to divide those counts by the total number of mapped reads in each sample in millions, followed by the size of each transcript in kb to obtain RPKM values. A previously published housekeeping gene, human C1orf43, (Eisenberg and Levanon 2013) was selected as a control (Supplementary Fig. S13), and orthologs in other species were identified using Ensembl gene trees (Zerbino, et al. 2018).

To search for stem loop sequences and poly(A) signals, we first extracted 600bp of genomic sequence on each side of the stop codon of each H2B variant. We downloaded a model for the histone 3’ UTR stem loop (accession RF00032) from the RFAM database (Kalvari, et al. 2021) searched for matches using the ‘cmsearch’ algorithm (covariance model search) of the Infernal package (Nawrocki and Eddy 2013). For our analysis of poly(A) signals, we searched for exact matches to AATAAA or ATTAAA, the two most commonly-found signal sequences in human transcripts (Beaudoing, et al. 2000). This approach is somewhat limited, however. These short motifs will yield many false positive matches, and previous analysis shows that many polyadenylated human transcripts have no recognizable signal sequence (Beaudoing, et al. 2000).

## Data access

Newly annotated sequences identified in this study have been submitted to GenBank (https://www.ncbi.nlm.nih.gov/genbank/).

## Acknowledgments

We thank Ching-Ho Chang, Courtney Schroeder, Sierra Simmerman, and Paul Talbert for comments on the manuscript. This work was supported by grants from the NIH (National Institute of General Medical Sciences) R01 GM074108 and from the Howard Hughes Medical Institute to H.S.M. The funders played no role in study design, data collection and interpretation, or the decision to publish this study. H.S.M. is an Investigator of the Howard Hughes Medical Institute.

