## Supplementary Figures 1-14 for "Novel classes and evolutionary turnover of histone H2B variants in the mammalian germline"

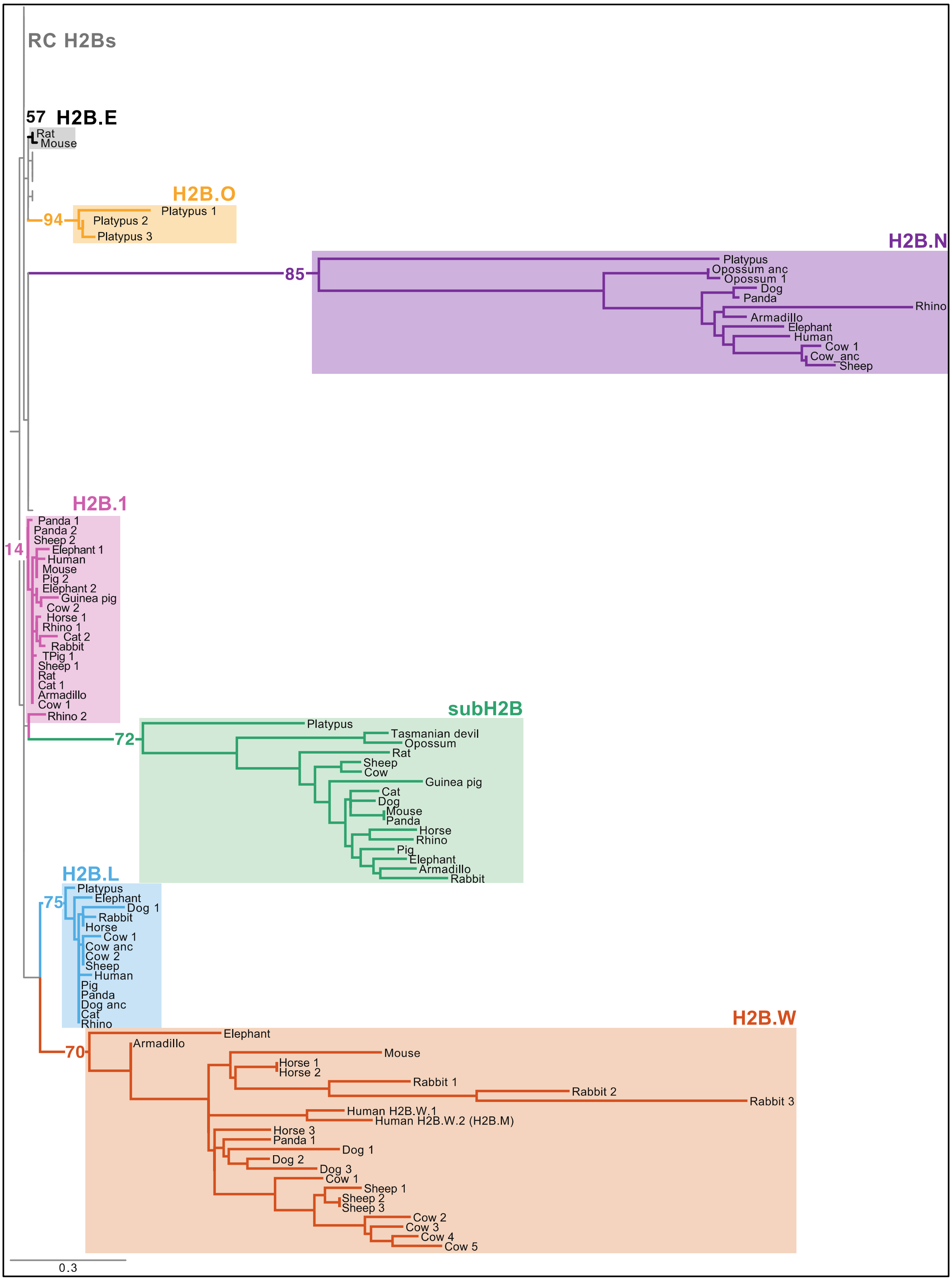


**Supplementary Figure 1. Phylogeny of mammalian H2B variants using the histone fold domain and αC domain.**
Maximum-likelihood protein phylogenetic tree of the histone fold domain (HFD) and αC domain of selected RC H2Bs and all identified H2B variants from eighteen representative

mammalian species (same as Figure 1A, Supplementary Data S1). Bootstrap values at selected nodes with >50% support are shown. Asterisk (*) indicates low bootstrap support (14%) for H2B.1 likely due to high conservation of HFD between H2B and H2B.1 (also see Supplementary Figure S5). Eight H2B variants identified are highlighted in colored boxes: H2B.E (Black), H2B.O (yellow), H2B.N (purple), H2B.1 (pink), subH2B (green), H2B.L (blue), and H2B.W (orange). Names of mammals are indicated at branch tips. In species with multiple copies of a variant, ancestral copies identified based on syntenic location are indicated with '_anc', and other duplicates are indicated with numbers (*e.g.,* Cow 1 in H2B.N group).

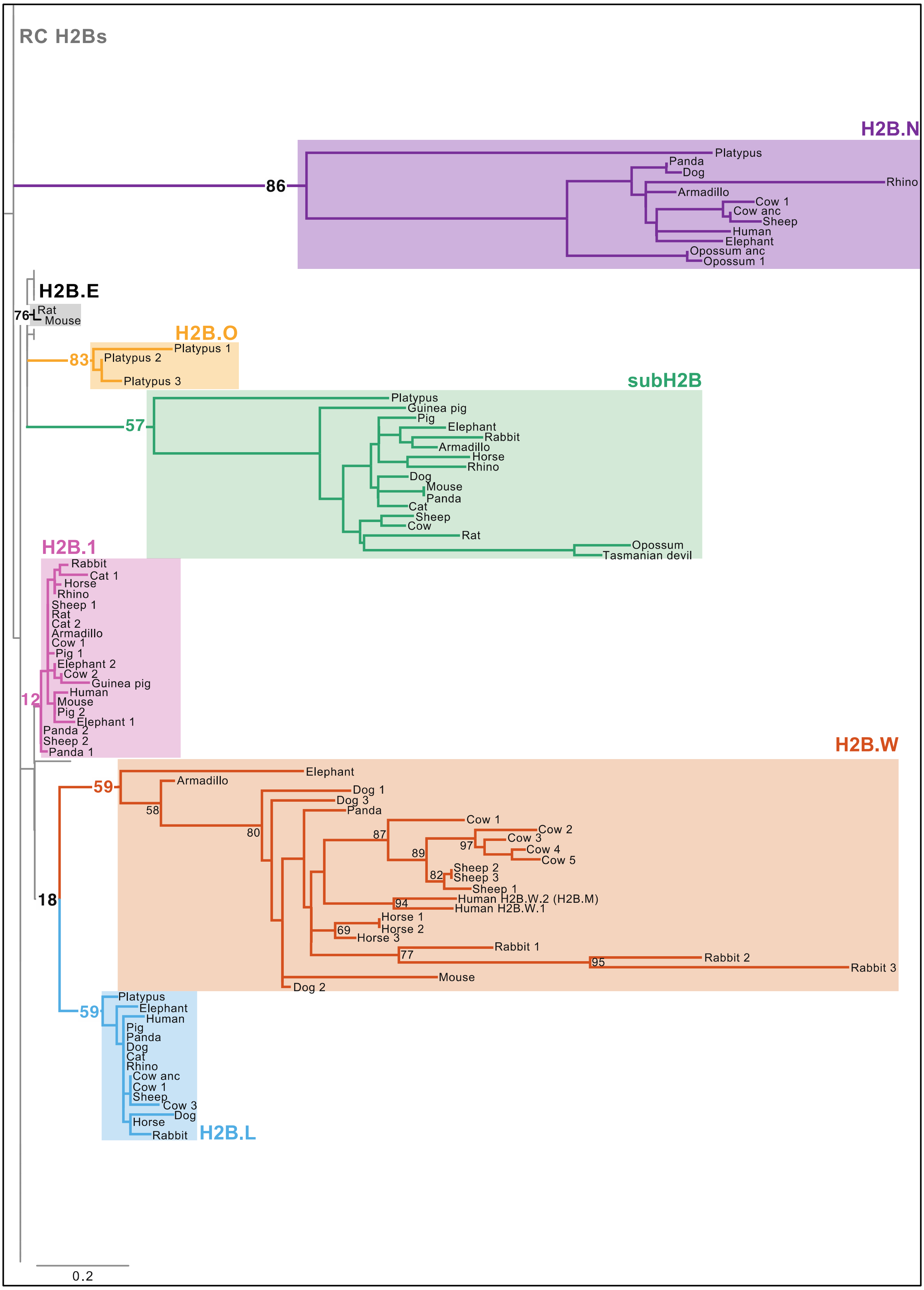


**Supplementary Figure 2. Mammalian H2B phylogeny using the histone fold domain.** Maximum-likelihood protein phylogenetic tree of the histone fold domain (HFD) of RC H2Bs and H2B variants from eighteen representative mammalian species (same proteins as in

Supplementary Figure 1). Bootstrap values at selected nodes with >50% support are shown. Eight H2B variants identified are highlighted in colored boxes as in Supplementary Figure 1.

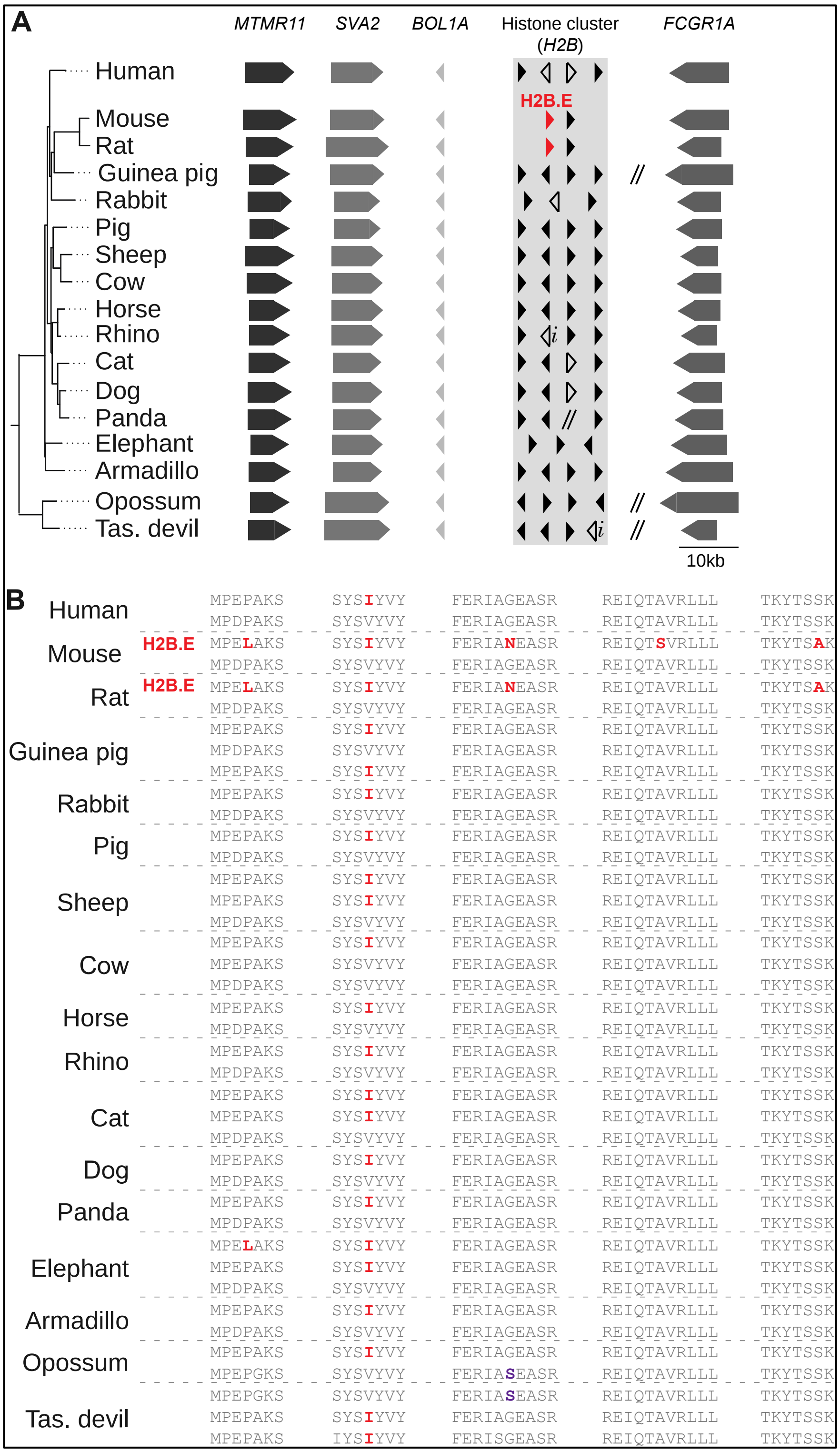


**Supplementary Figure 3. Synteny and sequence of H2B.E.**

(A) All H2Bs (H2B cluster, black triangles) found within the syntenic locus of histone variant H2B.E in representative mammalian species are shown. For simplicity, H2A, H3 and H4 variants also present at this location are not shown. Double slashes indicate breaks in synteny. Within the H2B cluster, *i* indicates incomplete genome information and empty triangles indicate inferred pseudogenes.

(B) Alignment of regions surrounding H2B.E diagnostic residues (red) of unique H2B sequences within the histone cluster in (A) are shown. Diagnostic residues in opossum and

Tasmanian devil in one copy of H2B were different from both H2B.E and the remaining H2Bs (purple).

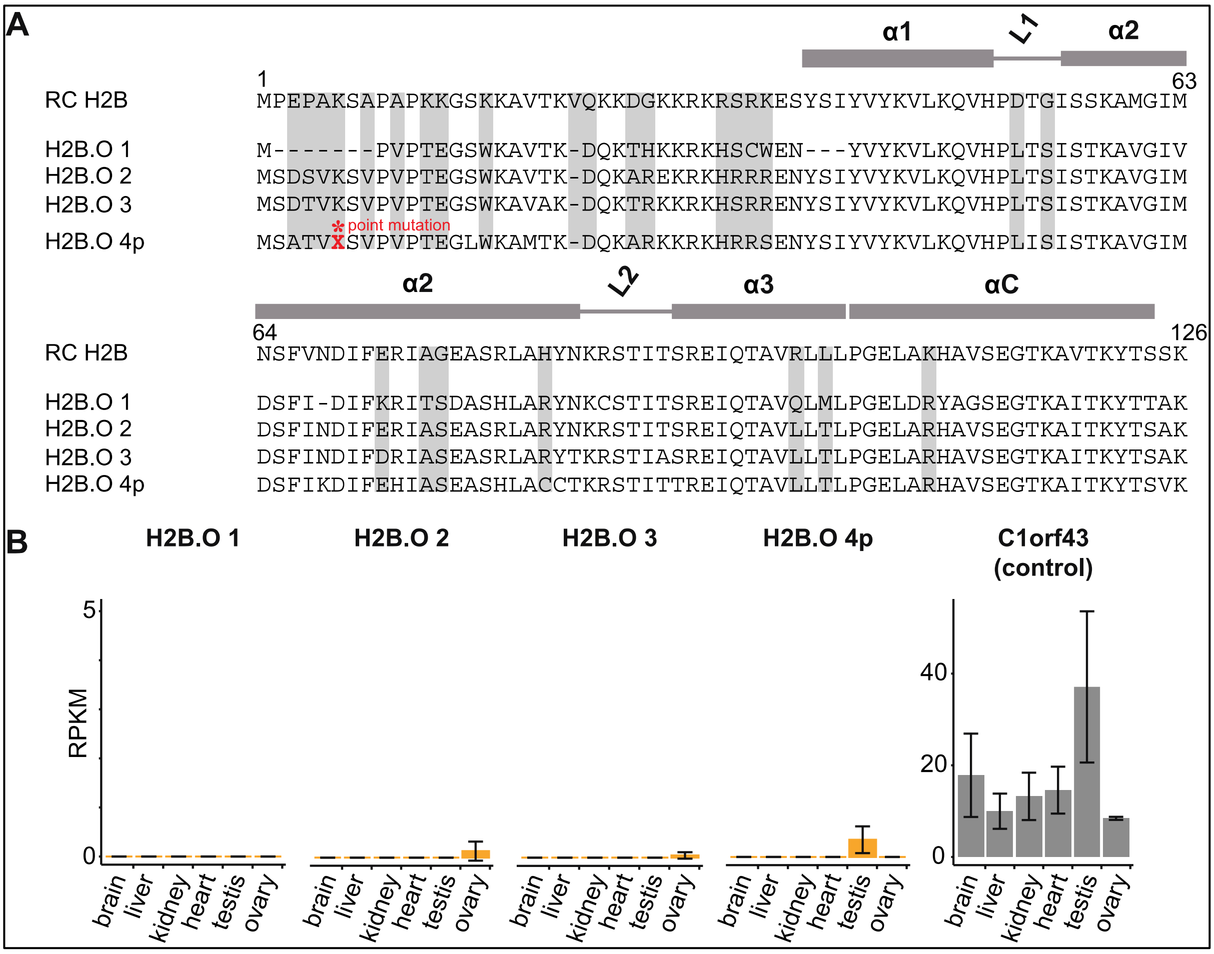


**Supplementary Figure 4. Alignment and expression of H2B.O in platypus.**

(A) Sequence alignment of RC H2B and platypus H2B.O variant residues. Residues that are different between RC H2B and a majority of H2B.O variants are highlighted in grey. A single nucleotide mutation that results in a premature STOP codon in the open reading frame (ORF) of H2B.O 4 variant is indicated with a red asterisk. This mutation could either suggest early pseudogenization of the variant or a sequencing error.

(B) RNA expression of H2B.O variants and a control gene, *C1orf43* (reads per kilobase per million mapped reads, RPKM) in publicly available bulk RNA-seq data across somatic and germline tissues in platypus are plotted.

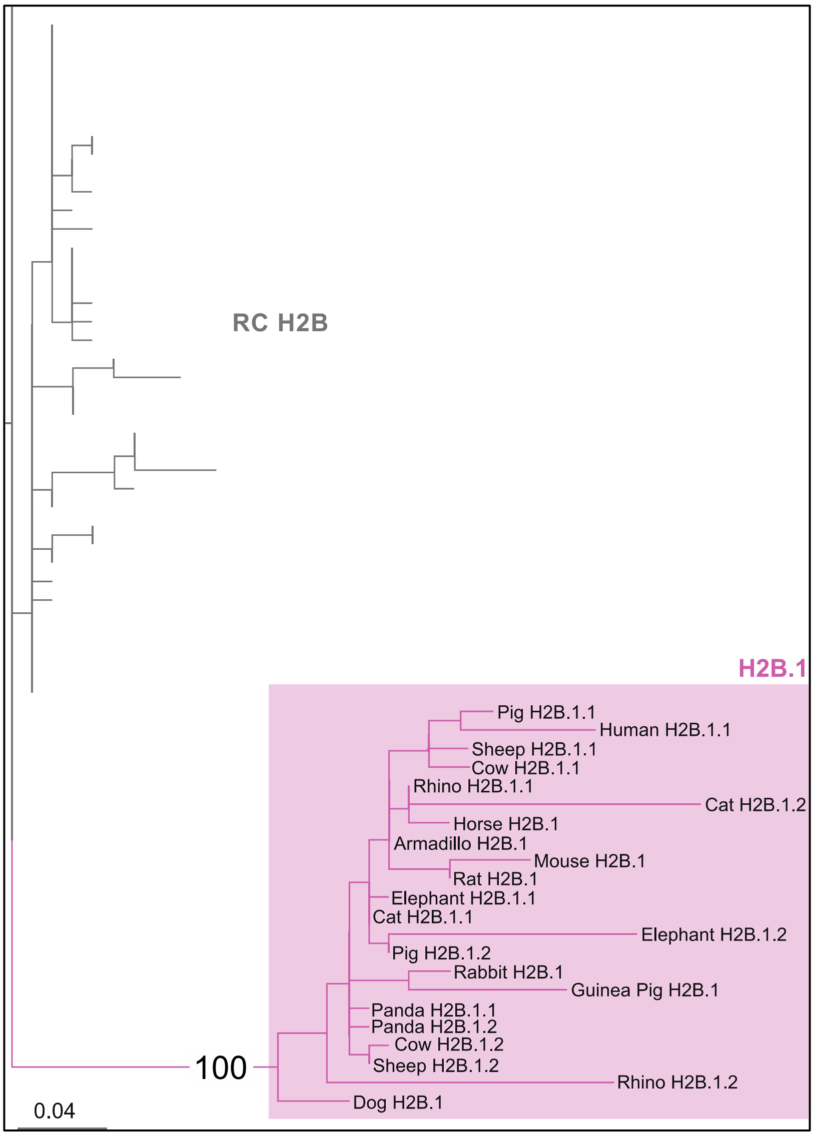


**Supplementary Figure 5. Phylogeny of full-length mammalian H2B.1 and RC H2B.** Maximum-likelihood protein phylogenetic tree of the full-length protein sequence of selected RC H2B sequences and the H2B.1 variant (pink box) from eighteen representative mammalian species (Supplementary data S1). Bootstrap values for H2B.1 ancestral node is shown. Duplicates of H2B.1 were found at the same syntenic location in some mammals (Supplementary Figure S4) and are indicated with numbers.

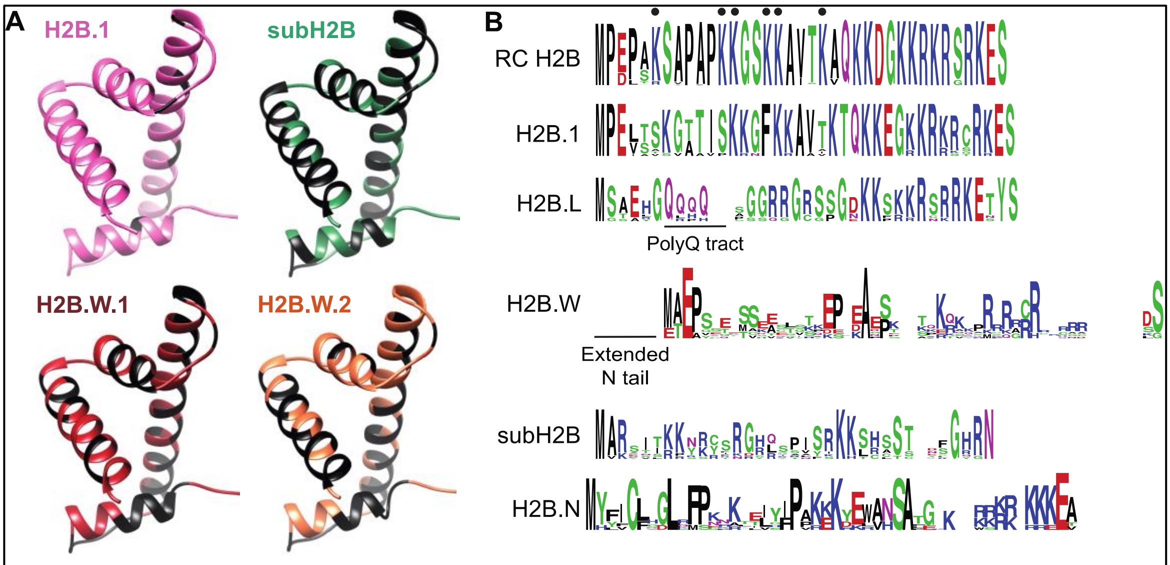


C

| Human Histone | Full length | | HFD | |
|---|---|---|---|---|
| | Charge | pI | Charge | pI |
| RC H2B | +17.33 | 10.16 | +4.3 | 9.47 |
| H2B.1 | +17.33 | 10.16 | +4.3 | 9.47 |
| H2B.L | +12.08 | 9.94 | +2.14 | 8.85 |
| H2B.W.1 | +12.17 | 10.48 | +4.39 | 10.38 |
| H2B.W.2 | +8.2 | 9.93 | +2.39 | 9.51 |
| H2B.N | +17.23 | 10.4 | +5.27 | 9.99 |

**Supplementary Figure 6. Characteristics of H2B variants.**

(A) Homology models of human H2B.1, rhesus macaque subH2B, human H2B.W.1 and human H2B.W.2 with sites that differ from RC H2B are highlighted in black (see Methods).

(B) Logo plots depicting protein alignments of the N terminus of RC H2B and H2B variants across an identical set of representative mammals (see methods). Color of residues highlight biochemical properties: hydrophobic (black), positively charged (blue), negatively charged (red), polar (green) and others (purple). Above the RC H2B plot we highlight residues that are post-translationally modified (filled circles). In some species, H2B.L has an extended poly Q tract. The H2B. W clade has a poorly-conserved extended N terminal tail.

(C) The isoelectric points (pI) and charges of human H2Bs (full-length protein and histone fold domain).

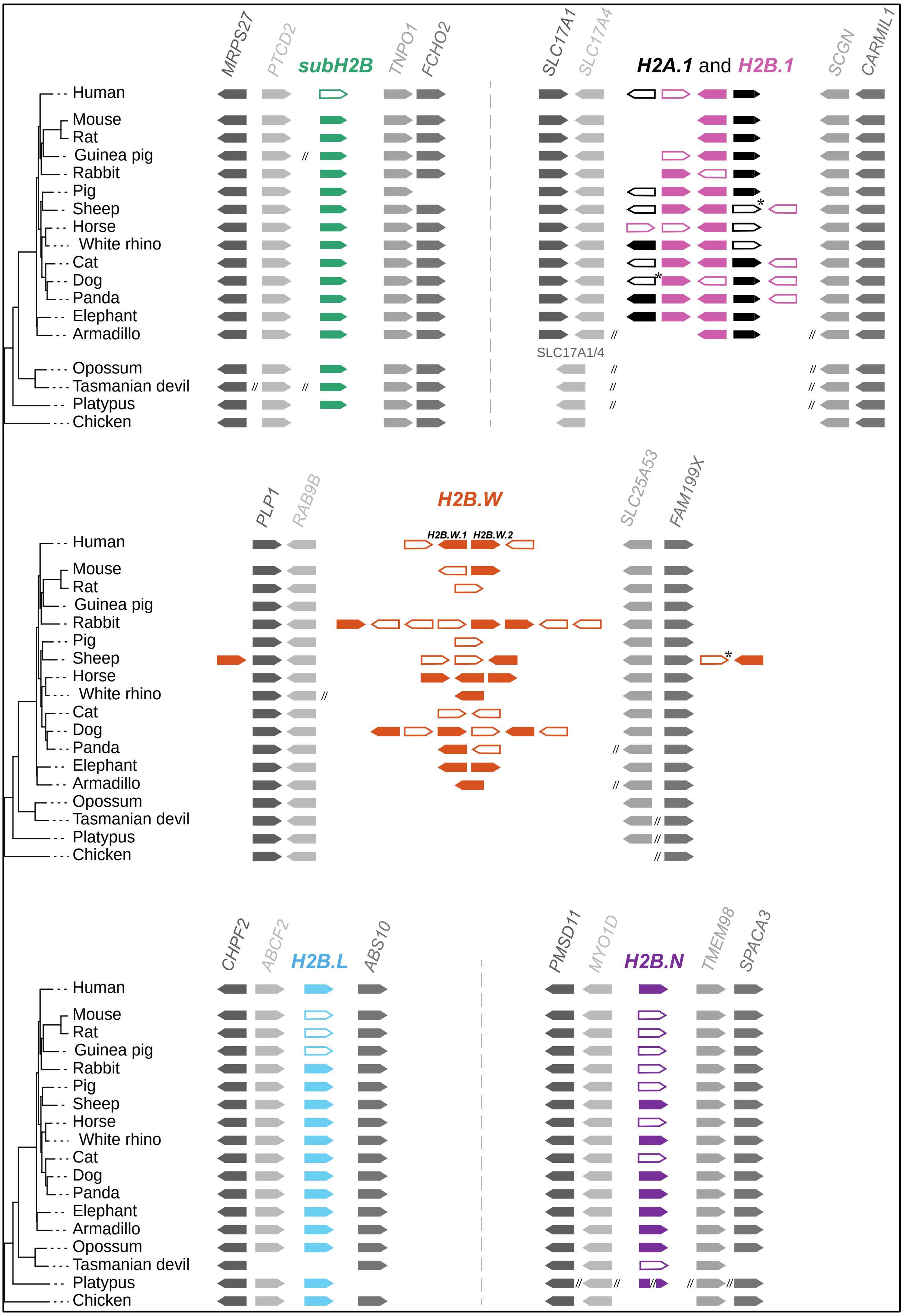


**Supplementary Figure 7. Synteny of all germline-specific mammalian H2B variants.** Syntenic location and retention of H2B variants in mammals and a non-mammalian outgroup, chicken. Each gene is represented as an arrow to indicate gene orientation.

Variants are indicated in colors as in Figure 1 and conserved neighbouring genes are indicated in shades of grey. Double slashes indicate breaks in synteny. Inferred pseudogenes are represented as empty boxes. Asterisks (*) indicate pseudogenization by a single nucleotide change which could either be sequencing error or a true mutation. Absence of an arrow indicates that the gene was not found– we did not distinguish true gene loss from absence because of gaps in genome assembly.

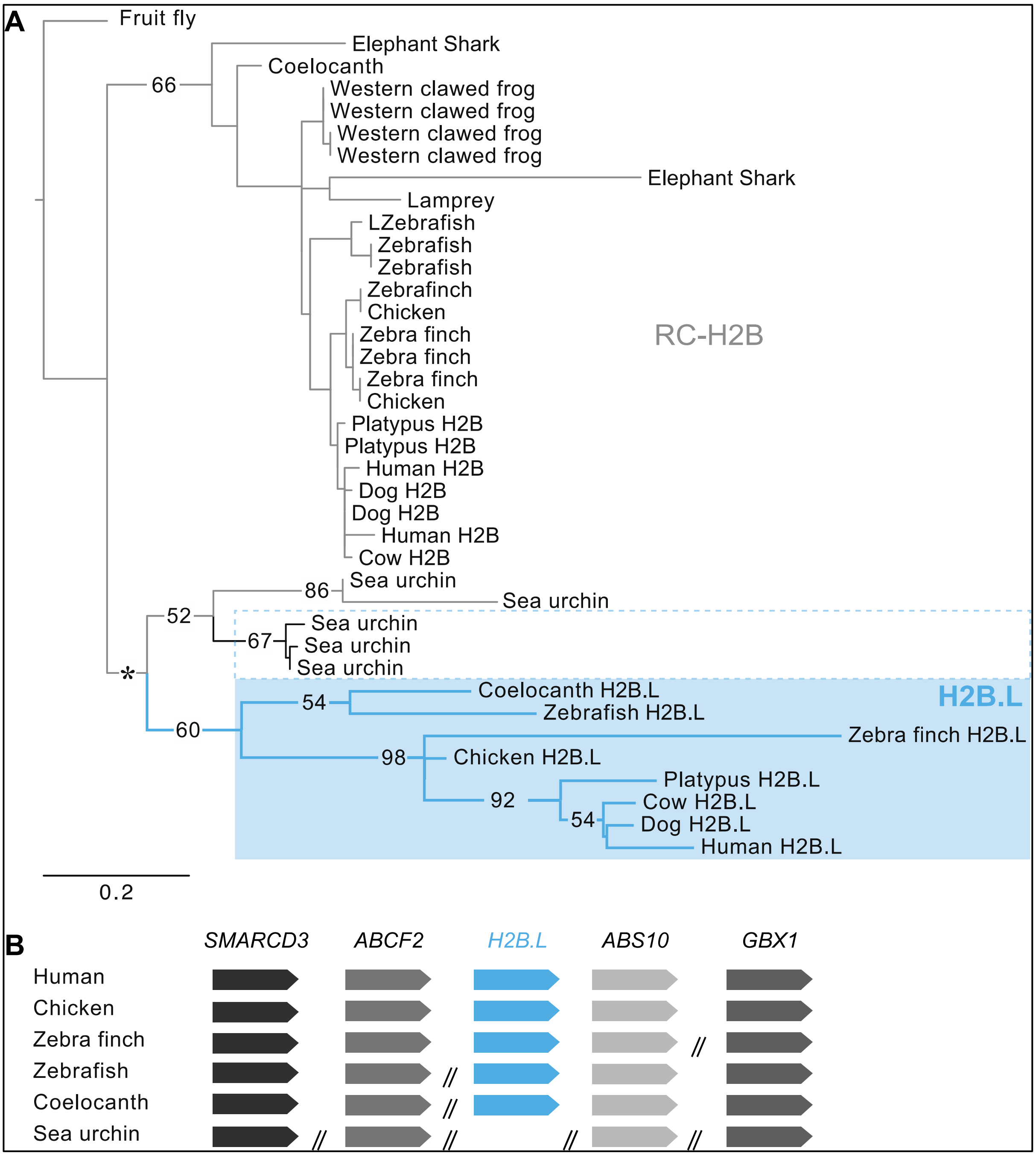


**Supplementary Figure 8. Phylogeny and synteny of H2B.L in vertebrates.**
(A) Maximum-likelihood protein phylogenetic tree of the full-length protein sequence of selected RC H2B and variant H2B.L (blue box) from representative vertebrate species (Supplementary Data S2). Bootstrap values at select nodes with >50% support are shown. Asterisk (*) indicates lower bootstrap support (45%). Late-histone H2B sequences in sea urchin (Lai et al., 1986, Kemler et al., 1986, Marzluff et al., 2006) are encompassed in a dashed box.
(B) Syntenic location of non-mammalian vertebrate H2B variants that group with mammalian H2B.L in (A) compared to human H2B.L synteny are shown.

```
                          1                                                               65
Rhesus macaque            MARSSTKKHKYSKRHQSPTSRKKA---HSSIDFVHGNYSFFVNKVLKEVVSHRGTSSRTLDLMNTLIN

Pseudogenized subH2B
Human                     MARSSTKKYKYSKRHQTPTSRKKAKKAYSSIDFGHGNYSFFVNKVLKEVVPCRGISSRTLDLMNALIN
Chimpanzee                MARSSTKKYKYSKRHQTPTSRKKAKKAYSSIDFGHGNYSFFVNKVLKEVVPCRGISSRTLDLMNALIN
Bonobo                    MARSSTKKYKYSKRHQTPTSRKKAKKAYSSIDFGHGNYSFFVNKVLKEVVPCRGISSRTLDLMNALIN
Gorilla                   MARSSTKKYKYSKRHQTPTSRKKAKKAYSSIDFGHGNYSFFVNKVLKEVVPCRGISSRTLDLMNALIN

                                                                  * point mutation
Gelada                    MARSSTKKHKYSKRHQSPTSRKKA---HSSIDFVHGNYXFFINKVLKEVVSHRGTSSCTLDLMNTLIN
Olive baboon              MARSSTKKHKYSKRHQSPTSRKKA---HSSIDFVHGNYXFFINKVLKEVVSHRGTSSRTLDLMNTLIN
Hamadryas baboon          MARSSTKKHKYSKRHQSPTSRKKA---HSSIDFVHGNYXFFINKVLKEVVSHRGTSSRTLDLMNTLIN
Sooty mangabey            MARSSTKKHKYSKRHQSPTSRKKA---HSSIDFVHGNYXFFINKVLKEVVSHRGTSSRTLDLMNTLIN
Drill                     MARSNTKKHKYSKRHQSPTSRKKA---HSSIDFVHGNYXFFINKVLKEVVSHRGTSSRTLDRMNTLIN
                          *point mutation
African green monkey      ITSSSTKKHKYSKRHQSPTSRKKA---HSSIDFVHGNYSFFVNKVLKEVVSHRGTSSRTLDLINTLIN
                          *point mutation
Proboscis monkey          TARSSTKKYKYSKRHQSPTSRKKA---HSSIDFVHGNYSFLVNKVLKEVVSHRGISSRTLDLMNTLIN

Golden snub-nosed monkey  MARSSTKKYKYSKRHQSPTSRKKA---HSSIDFVHGNYSFLVNKVLKEVVSHRGISSRTLDLMNTLIN
Black snub-nosed monkey   MARSSTKKYKYSKRHQSPTSRKKA---HSSIDFVHGNYSFLVNKVLKEVVSHRGISSRTLDLMNTLIN

Marmoset                  MAKSRTKKHKYSKRHQIPTSRKKAKKAHSSIDFGHRNYSFGINRVLKEVVPRRGISSRTLYLMNALIN


                          66                                                       122
Rhesus macaque            NFFQHISMKAYRLMYFRNCCTLTPEDILKAAYLLLPQKTANYAVAFGSEVFRRYVHS

                                          1nt deletion
Pseudogenized subH2B                           ⊥
Human                     FLFQHIAIKAYRLMYSRNRCTLPLKISXRQCICCCLRKQLTMQRLLEVKWSTDMST
Chimpanzee                FLFQHIVIKAYRLMYSRNRCTLPLKISXRQCICCCLRKQLTMQRLLEVTWSTDMSTP
Bonobo                    FLFQHIVIKAYRLMYSRNRCTLPLKISXRQCICCCLRKQLTMQRLLEVTWSTDMSIP
Gorilla                   FLFQHIAIKAYRLMYSRNRCSLPLKISXRQCICCCLRKQLTMQRLLEVKWSTDMSTP

Gelada                    NFFQHIAMKAYRLMYFRNRCTLTPEDILKAVYLLLPQKTANYAVAFGSEVFRRYVHS
Olive baboon              NFFQHIAMKAYRLMYFRNRCTLTPEDILKAVYLLLPQKTANYAVAFGSEVFRRYVHS
Hamadryas baboon          NFFQHIAMKAYRLMYFRNRCTLTPEDILKAVYLLLPQKTANYAVAFGSEVFRRYVHS
Sooty mangabey            NFFQHIAMKAYRLMYFRNRCTLTPEDILKAVYLLLPQKTANYAVAFGSEVFRRYVHS
Drill                     NFFQHIAMKAYRLMYFRNRCTLTPEDILKAVYLLLPQKTANYAVAFGSEVFRRYVHS

African green monkey      NFFQHISMKAYRLMYFRNRCTLTPEDILKAVYLLLPQKTANYAVAFGSEVFHRYVHS

Proboscis monkey          NFFQHIAMKAYRLMYFRNRCTLTPEDILKAVYLLLPQKTANYAVAFGSEVFRRYVHS

                                                                                   *point mutation
Golden snub-nosed monkey  NFFQHIAMKAYRLMYFRNRCTLTPDDILKAVYLLLPQKTANYAVAFGSEVFRRYVHSK.....188
Black snub-nosed monkey   NFFQHIAMKAYRLMYFRNRCTLTPDDILKAVYLLLPQKTANYAVAFGSEVFRRYVHSK.....188
                                              1nt deletion
                                                   ⊥
Marmoset                  FIVQHIVMKAYRLTYIRNCCTLTPEDTXRQCICCCLRKQLTMQWFLEVKGVYRYVYS
```

**Supplementary Figure 9. Pseudogenization of subH2B in some primates.**
Primate subH2B sequences with disruptions to the open reading frame (ORF) are shown. Rhesus macaque which has an intact subH2B sequence is shown for comparison. Mutations that disrupt the ORF are shown in red. X indicates stop codon.

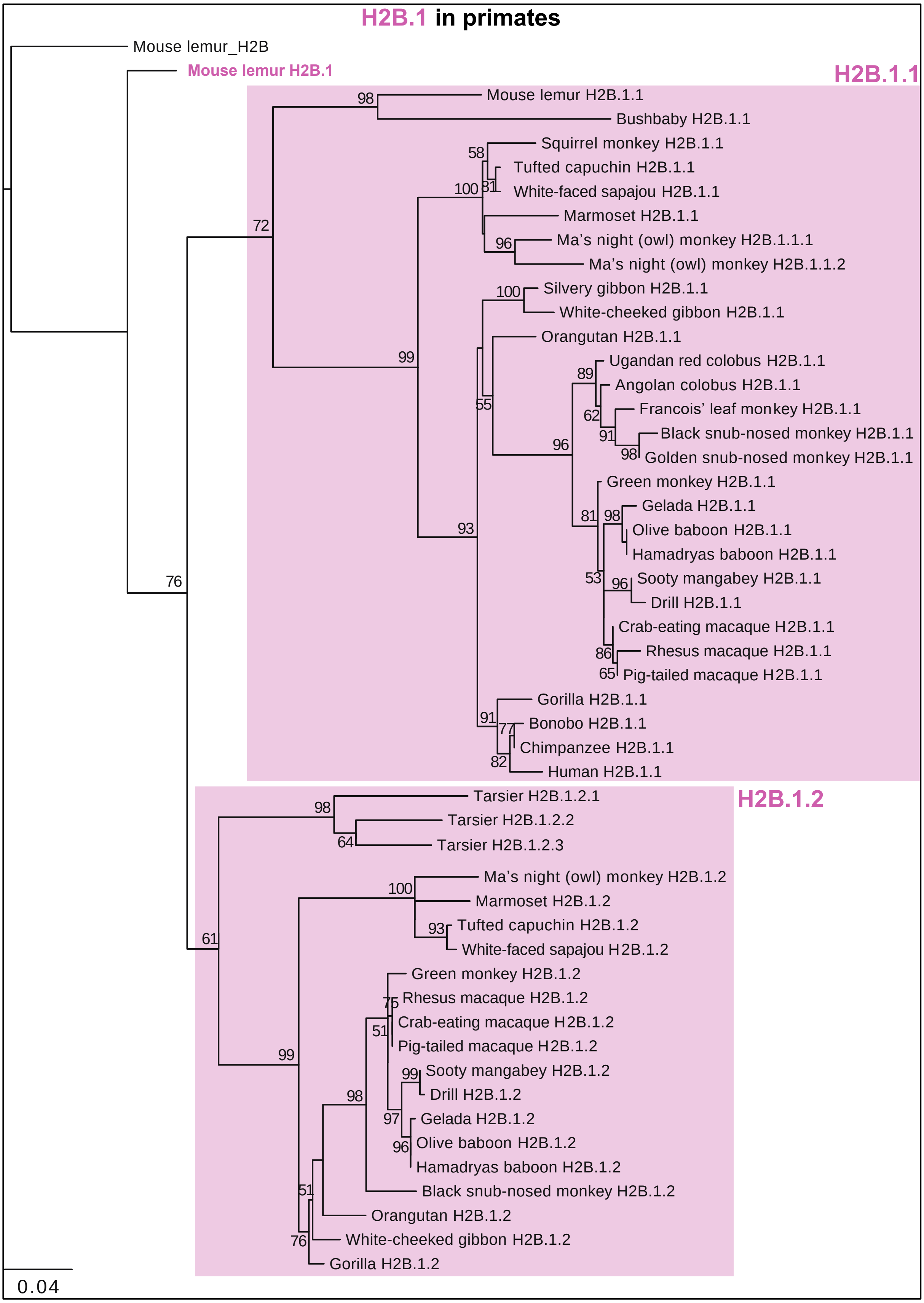


**Supplementary Figure 10. Phylogeny of H2B.1 in primates.**
Maximum-likelihood protein phylogenetic tree of the full-length protein sequence of mouse lemur RC H2B and variant H2B.1 (pink boxes) from thirty primate species (Supplementary

Data S3). Bootstrap values at all nodes with >50% bootstrap support are shown. All H2B.1 duplicates are local (i.e. found in the syntenic genomic locus) and are indicated with numbers at the end of gene names.

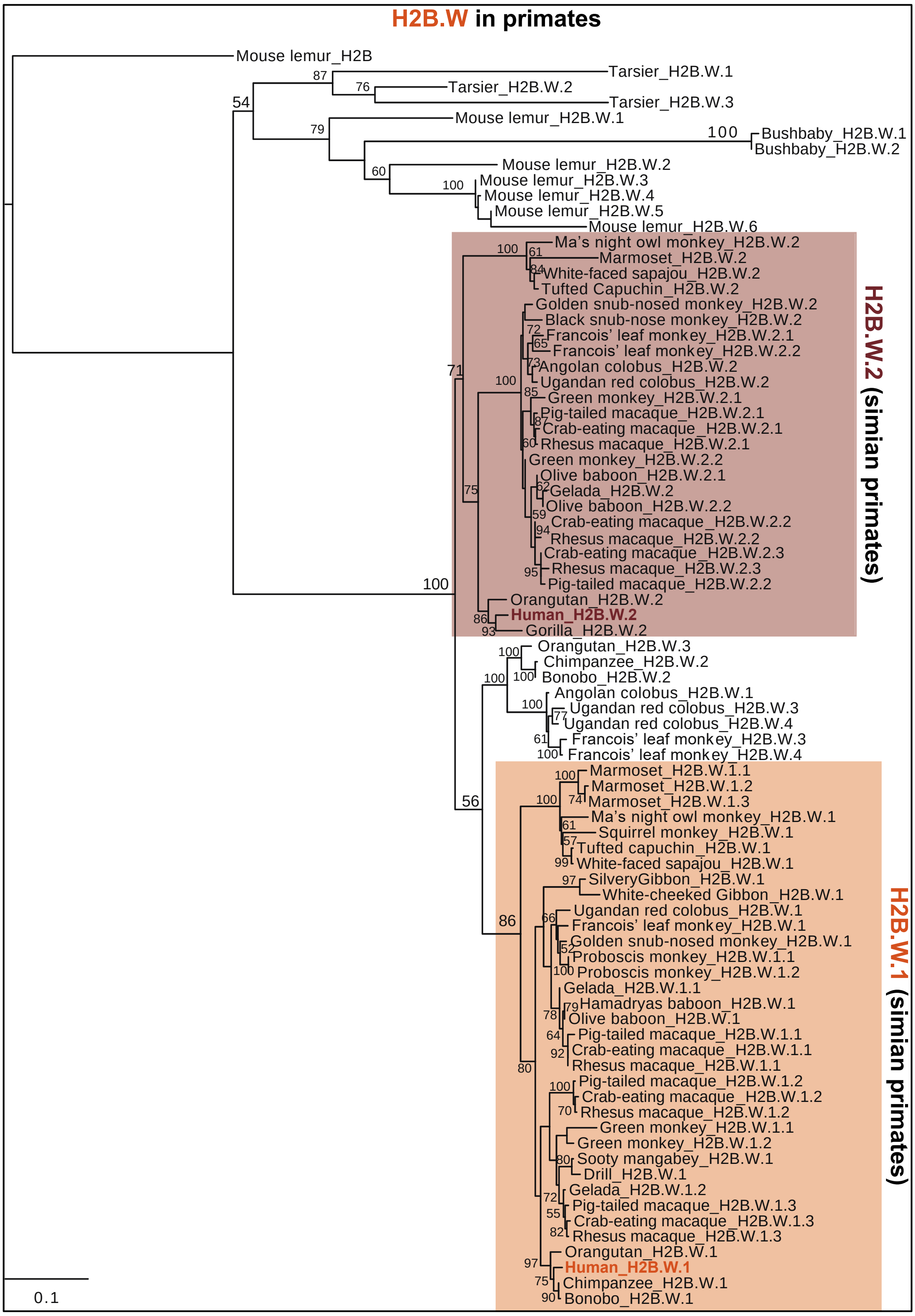


**Supplementary Figure 11. Phylogeny of H2B.W variants in primates.**
Maximum-likelihood protein phylogenetic tree of the full-length protein sequence of mouse lemur RC H2B and variant H2B.W.1 and H2B.W.2 from thirty primate species (Supplementary Data S4). Bootstrap values at all nodes with >50% bootstrap support are shown. All H2B.W duplicates are local (i.e. found in the syntenic genomic locus) and are indicated with numbers at the end of gene names.

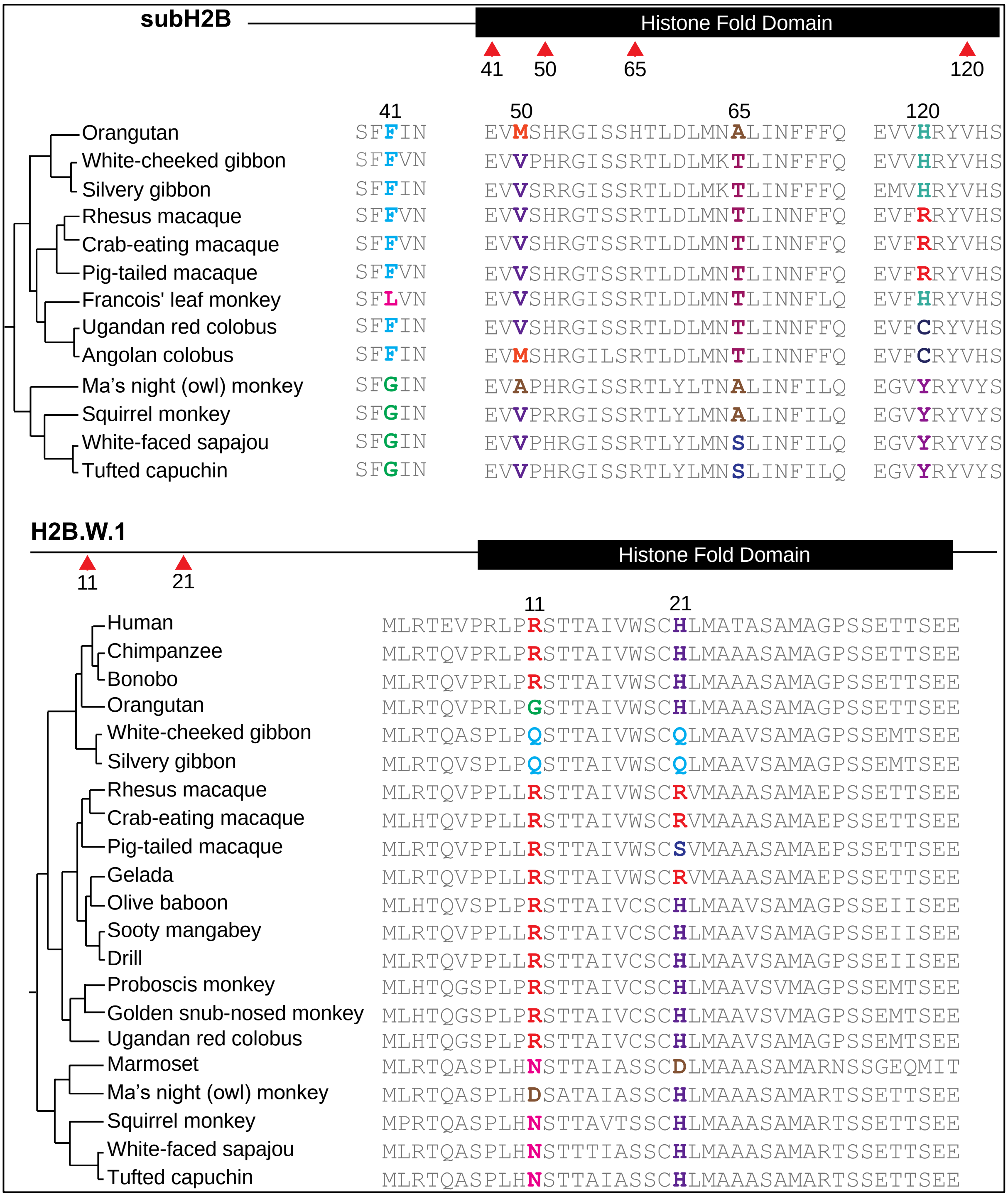


**Supplementary Figure 12. Positively selected residues of subH2B and H2B.W.** Alignments of regions surrounding positively selected sites (colored amino residues) in simian primate subH2B (top) and one copy of H2B.W.1 (bottom) genes identified by PAML. Red arrows indicate positions of positively selected residues on the protein schematic for each variant.

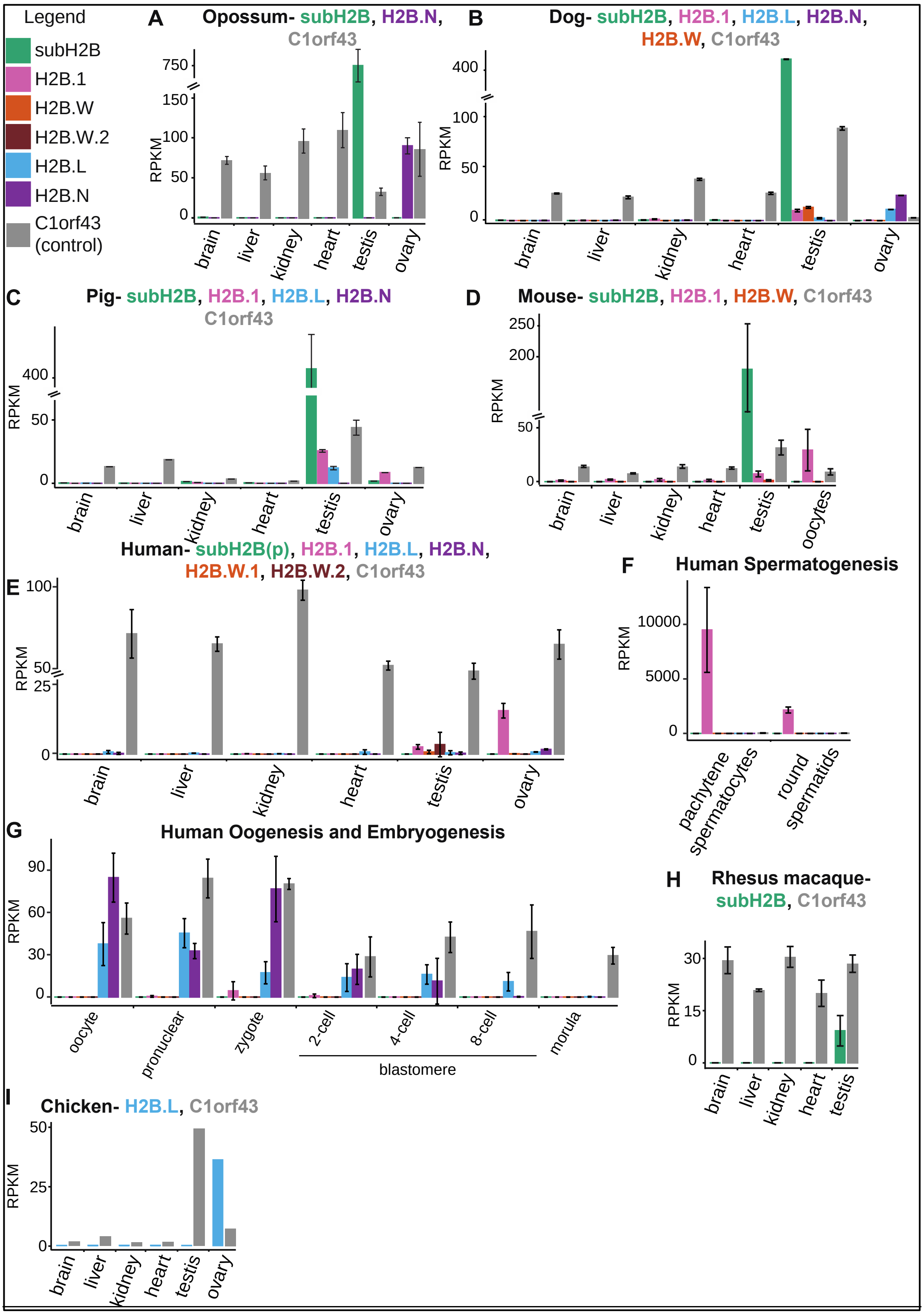


**Supplementary Figure 13. Expression of H2B variants in somatic and germline tissues of representative animals.**

Expression of select H2B variants in publicly available RNA-seq datasets from selected somatic tissues (brain, liver, kidney and heart) and reproductive tissues (testis and ovary) of (A) opossum, (B) dog, (C) pig, (D) mouse, (E-G) human (H) rhesus macaque and (I) chicken. Expression of variants that were inferred to be pseudogenes is not shown, except for human subH2B. Mapped reads are shown in reads per kilobase per million mapped (RPKM). The bar heights show median RPKMs of biological replicates and error bars show median absolute deviations, where available.

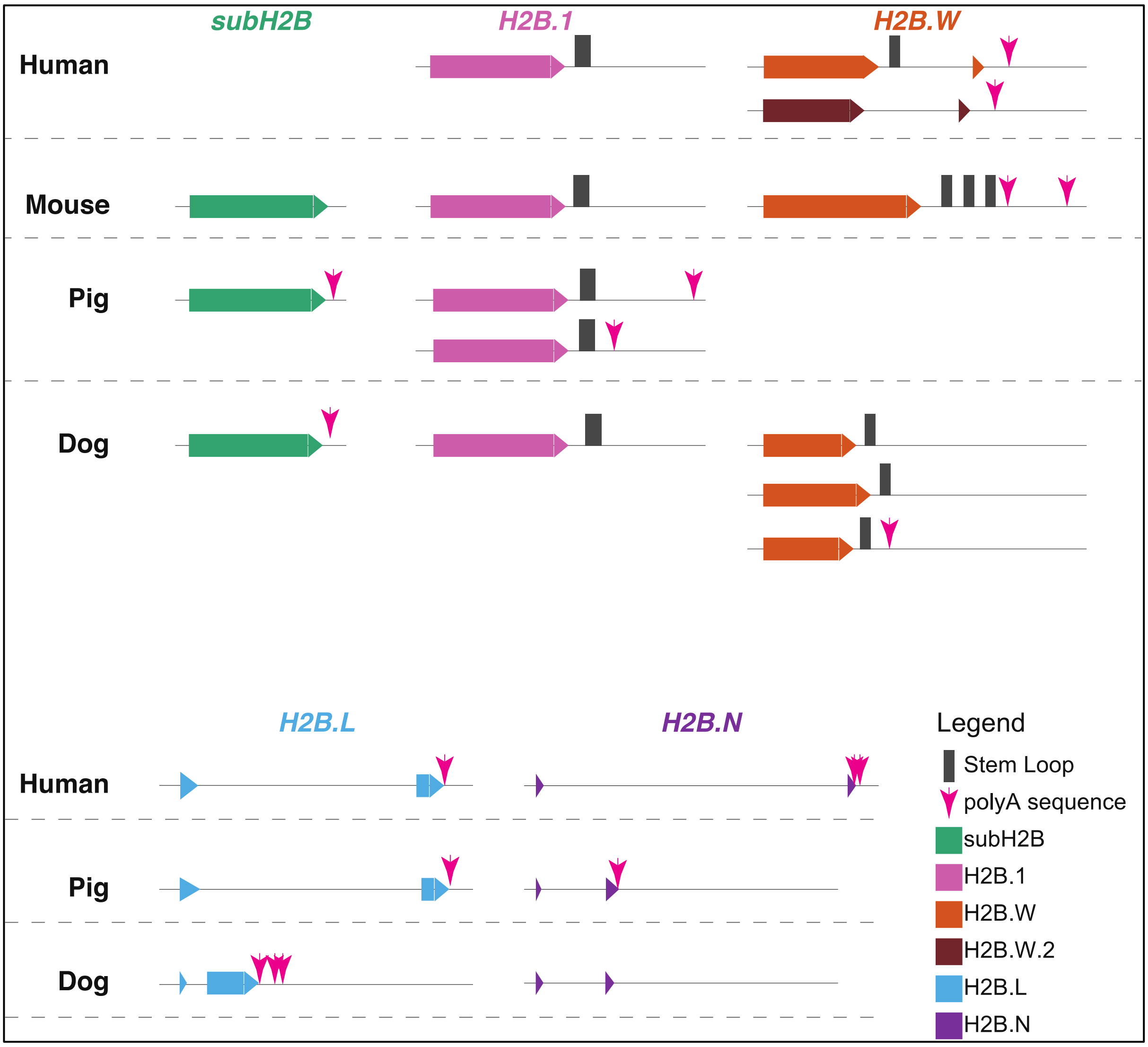


**Supplementary Figure 14. Detecting presence of poly(A) and stem loop structures in H2B variants of representative animals.**
Predicted stem loops (dark grey boxes) and poly(A) signals (pink arrows) are found in the 3' regions of H2B variant sequences of four representative mammals (human, mouse, pig and dog).
